# The impact of Parkinson’s disease on striatal network connectivity and cortico-striatal drive: an in-silico study

**DOI:** 10.1101/2023.09.15.557977

**Authors:** Ilaria Carannante, Martina Scolamiero, J. J. Johannes Hjorth, Alexander Kozlov, Bo Bekkouche, Lihao Guo, Arvind Kumar, Wojciech Chachólski, Jeanette Hellgren Kotaleski

**Affiliations:** Department of Computer Science, KTH Royal Institute of Technology, Stockholm, Sweden; Department of Mathematics, KTH Royal Institute of Technology, Stockholm, Sweden; Department of Neuroscience, Karolinska Institutet, Stockholm, Sweden

**Keywords:** Parkinson’s disease, Striatum, Computational modeling, Topological data analysis, Directed cliques, Network higher order connectivity, Neuronal degeneration model

## Abstract

Striatum, the input stage of the basal ganglia, is important for sensory-motor integration, initiation and selection of behaviour, as well as reward learning. Striatum receives glutamatergic inputs from mainly cortex and thalamus. In rodents, the striatal projection neurons (SPNs), giving rise to the direct and the indirect pathway (dSPNs and iSPNs, respectively), account for 95% of the neurons and the remaining 5% are GABAergic and cholinergic interneurons. Interneuron axon terminals as well as local dSPN and iSPN axon collaterals form an intricate striatal network. Following chronic dopamine depletion as in Parkinson’s disease (PD), both morphological and electrophysiological striatal neuronal features have been shown to be altered in rodent models. Our goal with this *in-silico* study is twofold: a) to predict and quantify how the intrastriatal network connectivity structure becomes altered as a consequence of the morphological changes reported at the single neuron level, and b) to investigate how the effective glutamatergic drive to the SPNs would need to be altered to account for the activity level seen in SPNs during PD. In summary we predict that the richness of the connectivity motifs in the striatal network is significantly decreased during PD, while at the same time a substantial enhancement of the effective glutamatergic drive to striatum is present.

**AUTHOR SUMMARY:** This *in-silico* study predicts the impact that the single cell neuronal morphological alterations will have on the striatal microcircuit connectivity. We find that the richness in the topological striatal motifs is significantly reduced in Parkinson’s disease, highlighting that just measuring the pairwise connectivity between neurons gives an incomplete description of network connectivity. Moreover, we predict how the resulting electrophysiological changes of SPN excitability together with their reduced number of dendritic branches affect their response to the glutamatergic drive from cortex and thalamus. We find that the effective glutamatergic drive is likely significantly increased in PD, in accordance with the hyperglutamatergic hypothesis.

## INTRODUCTION

Parkinson’s disease (PD) is a progressive neurodegenerative disease, debilitating both motor and cognitive systems. The progressive and chronic loss of dopamine results in a variety of changes in the ongoing and stimulus evoked activity in the striatum, the input stage of the basal ganglia (Figure 1A) (Sharott, Vinciati, Nakamura, and Magill (2017), Ketzef et al. (2017), Filipović et al. (2019)), globus pallidus (Mallet et al. (2008), Raz, Vaadia, and Bergman (2000), Tachibana, Iwamuro, Kita, Takada, and Nambu (2011)) and subthalamic nucleus (Bergman, Wichmann, Karmon, and DeLong (1994)) in both non-human primate and rodent models.

**Figure 1.**
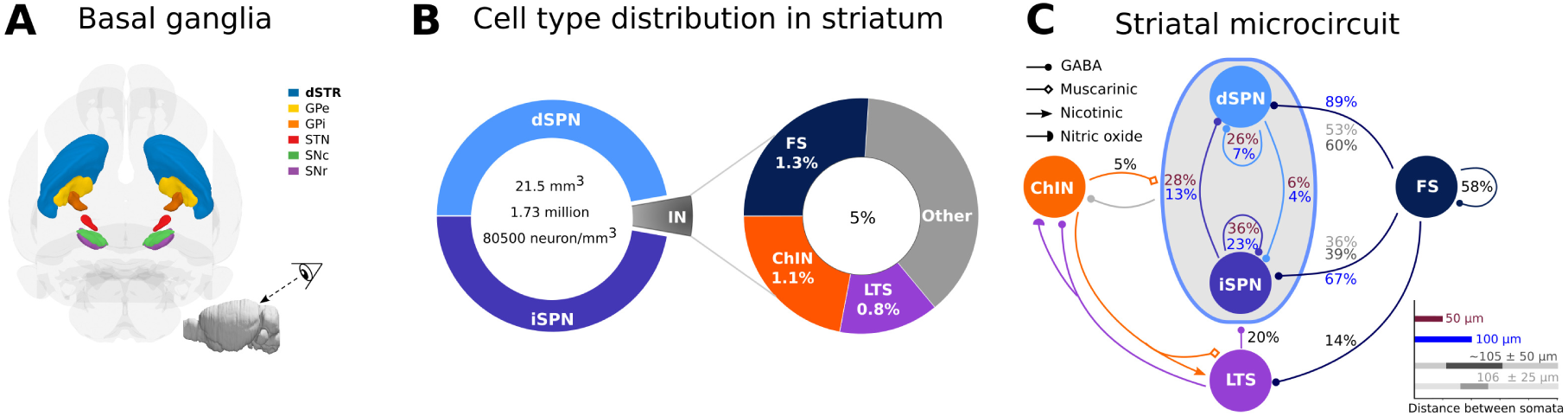
Organization of the striatal microcircuit. (A) View of the mouse basal ganglia nuclei (direction shown in the inset). The dorsal striatum (dSTR), globus pallidus external and internal segment (GPe and GPi, respectively), subthalamic nucleus (STN), substantia nigra pars reticulata and pars compacta (SNr and SNc, respectively) are shown in relative sizes. The colour coding is as indicated. The inset on the bottom right represents the entire mouse brain and the observer’s view. (B) The mouse striatum is around 21.5 mm^3^ with a total of 1.73 million neurons which correspond to approximately 80 500 neurons/mm^3^. The main cells in the striatum are the striatal projection neurons (SPN), they constitute around 95% of the neurons and they are divided into two subpopulations (dSPN and iSPN). The remaining 5% of the neurons are interneurons. Fast-spiking (FS), cholinergic (ChIN) and low-threshold spiking (LTS) interneurons are included in this *in-silico* network. Together they account for around 3.2% of the neurons (around 64% of the interneuron types). (C) Connection probabilities between the neuronal subtypes included in the *in-silico* network were collected from published data, as explained in Hjorth et al. (2020). When more than one number refers to the same connection (arrow) they come from different publications. In particular the distance between the somatic pairs is different. Dark brown and blue refer to somatic pair distance within 50 µm and 100 µm, respectively; while dark and light grey refer to an average distance of about 105 ± 50 µm and 106 ± 25 µm, respectively.

These neural activity alterations are accompanied by major changes in the morphology of the striatal projection neurons (SPNs). Postmortem analysis of neostriatal tissue reveals significant degeneration of the SPN dendrites in PD patients compared to controls without a history of neurological or neuropsychiatric illness. Neurodegeneration leads to a reduction by almost half of the total dendritic length and average length of the terminal dendritic segments at the most advanced stage of PD (McNeill, Brown, Rafols, and Shoulson (1988)). Dendritic degeneration is more pronounced in the putamen than in the caudate nucleus and is particularly dramatic in the commissural and post-commissural regions where the total dendritic length is reduced to less than a quarter (Zaja-Milatovic et al. (2005)).

Rodent models have been used to understand the striatal circuitry both in health and disease. SPNs account for about 95% of the striatal neurons in rodents and the remaining neurons are interneurons (Figure 1B) (Burke, Rotstein, and Alvarez (2017); Graveland and DiFiglia (1985)). SPNs are equally divided into two subpopulations expressing D1 or D2 dopamine receptors. The former, denoted dSPN, gives rise to the direct pathway and the latter, iSPN, the indirect pathway. In this computational study we have modelled the fast-spiking (FS), low threshold spiking (LTS) and cholinergic (ChIN) interneurons in addition to the two types of SPNs (as described in detail in Hjorth et al. (2020)). Some neurotoxin-induced and genetic rodent models of Parkinson’s disease are known to reproduce the dendritic degeneration of SPNs, but the reduction of total dendritic length is not as dramatic as in human patients at the terminal stages of PD (for example: 6-OHDA model in Fieblinger et al. (2014), Fieblinger et al. (2018); aphakia model in Alberquilla, Gonzalez-Granillo, Martín, and Moratalla (2020); knockout D1R mice (D1R*^−/−^*) in Suarez, Solis, Sanz-Magro, Alberquilla, and Moratalla (2020)). Loss of SPN dendrites reduces both the SPN intrastriatal connectivity (Taverna, Ilijic, and Surmeier (2008)) as well as the number of the glutamatergic synapses (extrastriatal connectivity) (Fieblinger et al. (2014), Zhai, Shen, Graves, and Surmeier (2019)). In contrast to SPNs, fast spiking interneurons (FS) may exhibit axonal sprouting (over 60% longer than in control) and formation of new functional FS synapses of similar strength specifically onto iSPNs (Gittis et al. (2011)). This reorganization of the local striatal network happens rapidly within the first week after the 6-OHDA lesion and precedes dendritic atrophy in SPNs.

Striatum is crucial for sensori-motor integration (Wall, De La Parra, Callaway, and Kreitzer (2013), de la Torre-Martinez, Ketzef, and Silberberg (2023)), action-selection (Redgrave, Prescott, and Gurney (1999)) and reinforcement learning (Doya (2008)). To better understand how these functions of the striatum are impaired by loss of dopamine it is important to characterise and predict how changes in the single neuron morphologies affect the network connectivity structure and representation of cortical and thalamic inputs.

To characterise the impact of the progressive loss of SPN dendrites and sprouting of FS axons on network connectivity we use the digital reconstruction of the mouse striatal microcircuitry as in Hjorth et al. (2020). Here, multicompartmental neuron models based on reconstructed morphology, ion channels expression (modelled using the Hodgkin-Huxley formalism) and electrophysiological *ex vivo* experimental rodents data were used (Hjorth et al. (2020)). Network connectivity is then generated based on touch detection between dendrites and axons combined with pruning rules to match experimental connection probabilities (Figure 1C), as described in previous studies of striatal (Hjorth et al. (2020)) and neocortical (Markram et al. (2015), Reimann, King, Muller, Ramaswamy, and Markram (2015)) microcircuitry. In this *in-silico* striatal microcircuit we systematically modify the neuron morphologies in order to match the single-cell morphometry observed in Parkinson’s disease and calculated not only the first order network properties (neuron degree and connection probability) but also quantified network connectivity motifs called **directed cliques**. Directed cliques are structural feedforward motifs of all-to-all connected neurons which were recently used to capture higher order interactions in somatosensory cortex structural networks (Reimann et al. (2017)). In this study we found that progressive dendritic degeneration dramatically affects statistics of directed clique counts particularly at the later PD stages. Our analysis showed that interneurons (FS, LTS and ChIN), despite only accounting for 5% of the neurons, are key to the formation of high dimensional directed cliques. These results suggest that interneurons play a crucial role in shaping the striatal network structure.

Next, to understand how altered dendritic morphology and membrane properties influence the transfer of cortical inputs to the striatum we activated dSPN and iSPN with simulated cortical inputs and compared the control and the PD case. The SPN model parameters for the PD case were tuned to reproduce the electrophysiological changes observed in Fieblinger et al. (2014). We found that SPN loss of dendritic branches and corresponding glutamatergic synapses, as seen in PD condition, severely reduced neuron sensitivity to input rates as well as correlations of the cortico-striatal input. To “compensate” for the loss of inputs, and investigate how to restore the SPN activity, we tested two strategies: a) strengthening of the remaining synaptic inputs, and b) rewiring by adding the lost glutamatergic synapses onto existing dendrites. Our results predict that at a single SPN level the effect of PD (i.e. the loss of dendrites and the corresponding synapses together with the altered membrane properties) can be counteracted by either rewiring or strengthening the cortico-striatal inputs. Moreover, SPN dendritic atrophy and sprouting of FS axons significantly depletes the richness of the striatal network connectivity in terms of number and size of higher order motifs. Loss of higher order striatal motifs highlights the importance of morphological changes in addition to changes in electrophysiological properties. While the activity of the single neurons (SPNs) can be restored by adjusting the synaptic inputs, the intrastriatal structural change resulting in the altered patterns of connectivity motifs would not be easily compensated for by simply increasing or decreasing the intrastriatal synaptic strengths. This is because directed cliques are formed based on the presence of intrastriatal synapses regardless of their strength, altering the strength does not affect clique formation. To induce changes in the number of higher order motifs only the rewiring strategy (and hence adding synapses) will be effective. Our work thus highlights the importance of being able to investigate separately the role of the structural and electrophysiological changes occurring in neurodegenerative diseases such as Parkinson’s disease. Here biophysically detailed *in-silico* reconstructions play an important role.

## RESULTS

PD progression is characterised by morphological changes of SPNs (dendritic atrophy) and FS (axonal growth) and changes in the neuron’s membrane properties. To disentangle such changes, we first investigate how the morphological alterations reshape the striatal circuitry. Subsequently we study how the loss of cortico-striatal synapses may affect the response of individual SPNs and potential compensatory mechanisms.

### Changes of single neuron morphology in PD striatum

We mimicked the gradual degeneration of the SPN (Fieblinger et al. (2014), Fieblinger et al. (2018), Alberquilla et al. (2020)) in PD by removing parts of the distal dendrites in three progressive stages (denoted PD1, PD2, PD3, see Materials and Methods). The control, non-PD stage, is denoted PD0. Examples of SPN PD morphologies, degenerated using the software ***treem*** (See Materials and Methods and Hjorth, Hellgren Kotaleski, and Kozlov (2021)), are illustrated in Figure 2 (A, dSPN, B iSPN, grey branches indicate the lost dendrites). Progressive degeneration of dendrites resulted in a reduction in the total dendritic length, number of branching points and number of primary dendrites consistent with the decrease reported in Fieblinger et al. (2014), Fieblinger et al. (2018), Alberquilla et al. (2020) (Figure 2C, (see Materials and Methods)).

**Figure 2.**
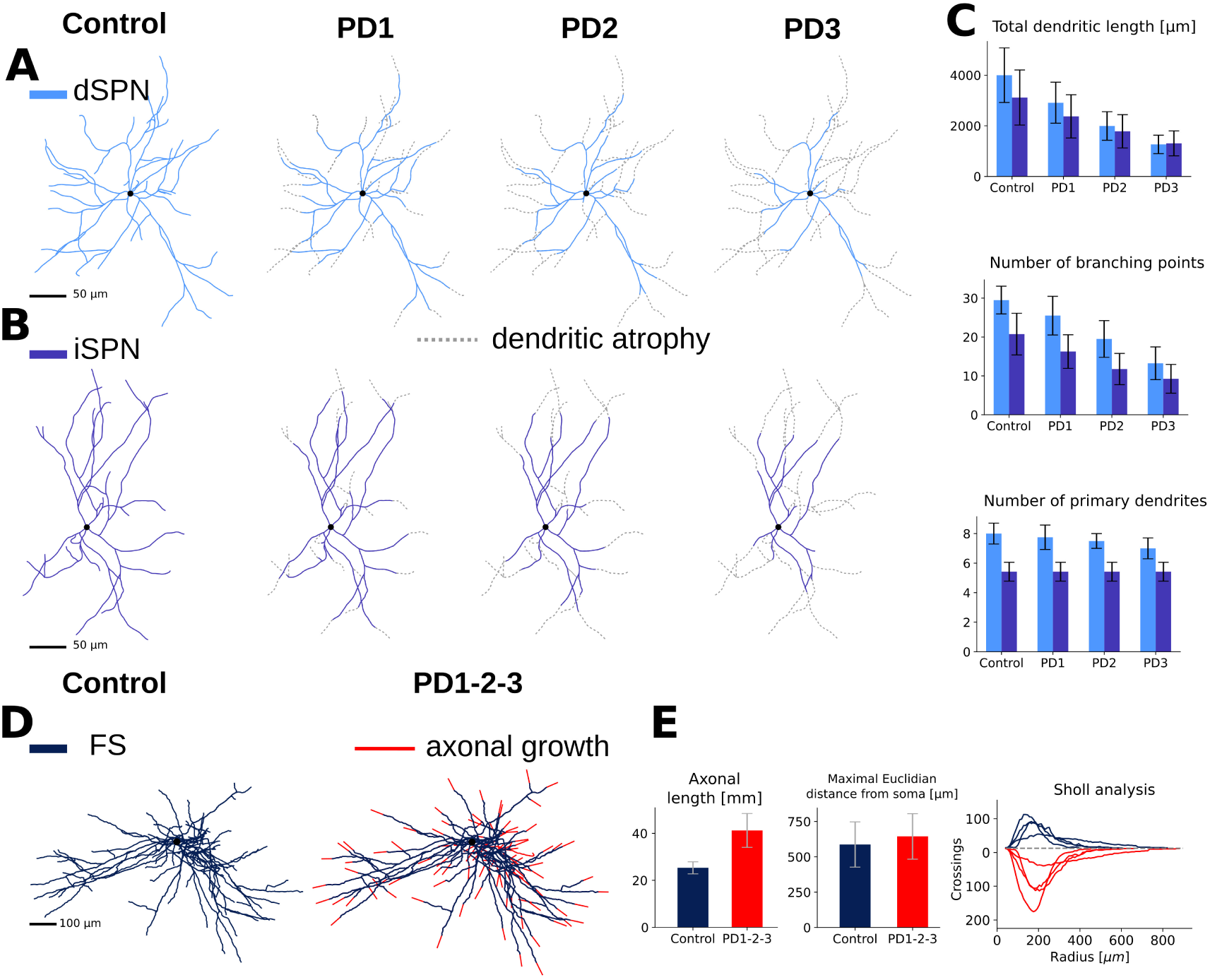
Morphological changes over Parkinson’s disease (PD) stages in the model. (A, B) The dendritic arborization of striatal projection neurons (dSPN and iSPN) is reduced in PD mice. Three different stages of the disease are simulated. PD1 refers to a mild starting phase, PD2 to a medium stage and PD3 to a very severe phase. Only soma and dendrites are shown for SPNs and the grey dotted lines represent the dendritic branches that atrophied. (C) Total dendritic length, number of branching points and number of primary dendrites for control (healthy) and PD stages are represented as histograms. (D) Axonal arborization of fast-spiking interneurons (FS) is increased in PD mice. Only soma and axon are shown for FS and the red lines represent the axonal branches that have sprouted. (E) The axonal length increased over 60% while the maximal euclidian distance from soma (the radius of the smallest sphere containing the axon) does not change significantly. The significant increase reported in the number of grid crossings by FS axons in PD (Gittis et al. (2011)) is also captured in our models and is shown in the Sholl analysis plot.

Next, as reported in Gittis et al. (2011), we modelled the increase in the FS axonal length using *treem* (Figure 2D, red branches indicate axonal sprouting; Figure 2E, left panel shows the increased total axonal length; for details see Materials and Methods). The average distance over which FS axons extended was not changed significantly in PD (Figure 2E, middle panel), but there was an increase in the number of grid crossings in the Sholl analysis (Figure 2E, right panel). In conclusion, the axonal trees of FS interneurons did not grow in a preferential direction but were denser than in control in accordance with Gittis et al. (2011).

### Predicting the network connectivity in the healthy and diseased state

Dendritic atrophy of SPN results in loss of both SPN local connectivity as well as decreased cortico-striatal and thalamo-striatal synaptic connectivity. On the other hand, FS axonal growth increases FS-iSPN connectivity and maintains almost invaried FS-dSPN connectivity. How these changes affect the striatal network structure beyond just a change in connection probabilities requires three dimensional reconstruction of the neuron morphologies and reconstruction of the network for different stages of PD progresion. To this end, we used the modelling framework ***Snudda***, presented in Hjorth et al. (2021, 2020). To create the reference microcircuit, we first randomly placed 100 000 neurons with appropriate cell densities in a cube (approximately 80 500 neurons/mm^3^) and then putative synapses between neuron pairs were detected based on proximity of their axons and dendrites. The initial touch detection overestimates the connectivity, therefore the putative synapses are pruned in successive steps (see Materials and Methods and Hjorth et al. (2021, 2020)) to match the experimental connection probability (Figure 1C). We refer to this as the healthy striatum or PD0. To avoid edge effects, when quantifying the local connectivity, only the very central neurons were considered for the analysis. In particular a subset of neurons closest to the centre of the cube and all their pre and postsynaptic neurons are selected (Figure 3 A, B, C; see Materials and Methods). We used the same strategy to generate the three stages of PD (see Materials and Methods). Because there is no cell loss, the distribution of the cells was retained (i.e. same as in the PD0 network) but due to SPN dendritic atrophy and sprouting of FS axons, synaptic connectivity becomes different (compare Figure 3 D, E and Supplementary Figure 1). During PD, because of the morphological alteration, the connection probabilities between cell types decreased for all connections except FS-iSPN (Figure 3F, G, H; Supplementary Figure 2).

**Figure 3.**
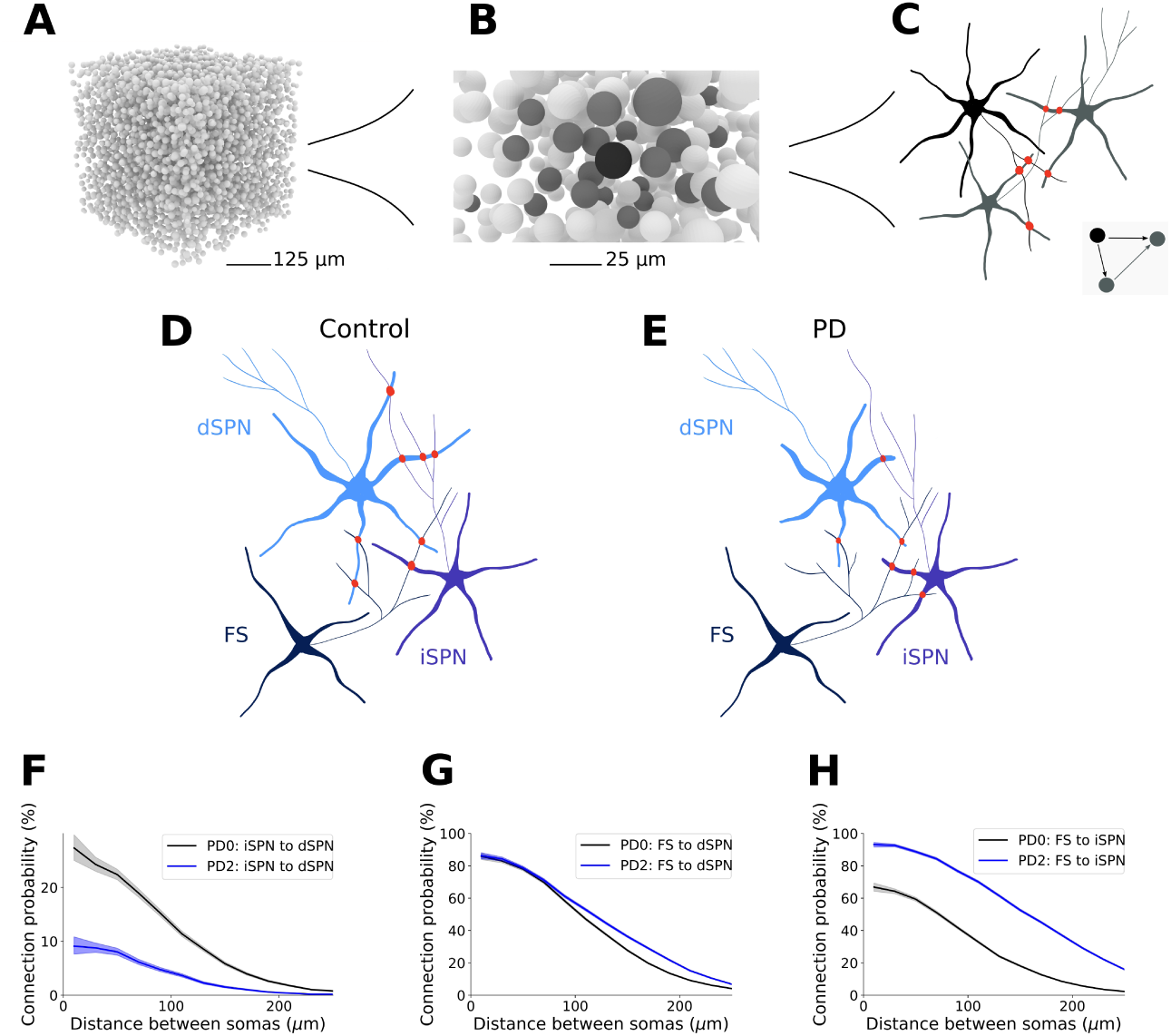
From morphologies to connectivity. Generating network connectivity using *Snudda* from reconstructed morphologies for healthy and PD networks. A) Example of positions of multi-compartmental neurons (somas) placed in a cube (5 000 somata are illustrated). A set of neurons in the centre of the cube, called kernel, is selected. All the pre- and post-synaptic neurons of the kernel form the core. The topological analysis is then performed on the kernel and core and only the cliques with at least one element in the kernel are kept to avoid edge effects. B) illustrates only one neuron forming the kernel (in black) and the elements in the core are in dark grey. C) Illustration of touch detection between a neuron in the kernel and two of its partners in the core. The three neurons together form a clique (inset figure) and the synapses are shown in red. (D, E) Illustration of connections between one FS, dSPN and iSPN in the PD0 network, and in PD2, respectively. The loss of dendrites in PD causes a reduction in connectivity between the two SPN neurons (here from 4 to 1), while the effect of FS axonal growth leads to new synapses on the iSPN (here from 1 to 3). F) The dendritic degeneration of the SPN leads to reduced pairwise connection probability at all soma-to-soma distances between SPN, here illustrated by the iSPN to dSPN connection probability. G) In accordance with data from Gittis et al. (2011) the growth of FS axons compensates for part of the degeneration of the dSPN morphologies, maintaining connection probability between the neuron types. H) For FS-iSPN connectivity the growth of the FS axons and locally increased synapse density compensates for the degeneration, leading to a doubling of the connectivity within 100 micrometres. Shaded regions in F, G and H represent the Wilson score interval.

### Topological characterization of the network in health and PD

#### Directed cliques in the healthy and PD models of striatum

A change in the pairwise connection probability is not informative about how the full connectivity has been restructured due to the modelled single-cell morphological changes (SPNs dendritic atrophy and FS axonal growth). To study the higher order properties of striatal networks we investigated the presence of specific motifs called directed cliques (Reimann et al. (2017)). A directed clique is a set of all-to-all connected neurons with a source and a sink. In this definition we are agnostic to the sign of the connection (excitatory or inhibitory). A directed clique constituted by *n* + 1 neurons is called a directed *n*-dimensional clique, or directed *n*-clique. For a rigorous definition of a directed *n*-clique see Materials and Method (Topological Measurements) and for examples or counterexamples of directed cliques see Figure 4 A1-4. These motifs are well suited to the study of the degeneration of striatal networks as in PD because they reveal complex patterns which are not visible from the analysis of single pairwise interactions.

**Figure 4.**
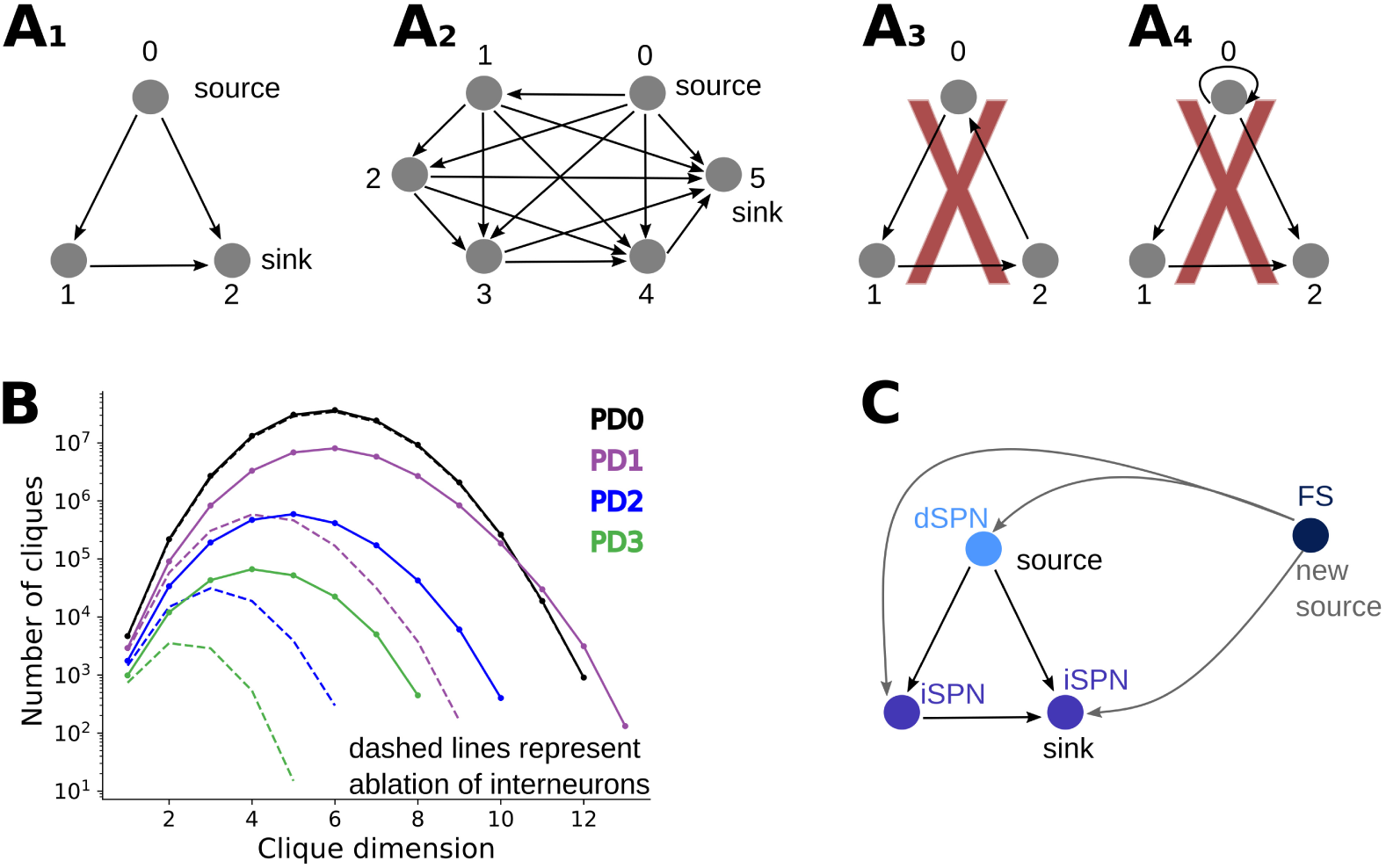
Directed cliques and their presence in PD networks. (A) A directed clique is a set of all to all connected vertices, with a unique source and a unique sink (see: Materials and Methods: Topological measurements). A clique composed of n+1 vertices is called a *n*-clique. (A1) and (A2) are examples of a 2-clique and a 5-clique respectively. Figure (A3) represents instead a cyclic structure in a directed graph, where a source and a sink are not present. This is therefore not an example of a directed clique. The graph represented in (A4) is also not a directed clique, according to our definition, since we assume that directed graphs do not contain self loops. (B) Number of directed cliques in the healthy network (PD0) and different Parkinsonian stages (PD1, PD2, PD3) as a function of the clique dimension in log-scale. Dashed lines represent directed clique counts in networks where interneurons have been ablated (only dSPNs and iSPNs are present in the networks). (C) Schematic representation suggesting how during PD stages new high dimensional cliques can be formed. The axonal growth of fast-spiking (FS) interneurons during PD progression can indeed determine connections from FS to existing directed cliques. The FS interneuron together with the neurons composing the already existing directed clique then form a new directed clique with source FS. This mechanism explains why PD1 has higher dimensional directed cliques than PD0, despite the dendritic atrophy of SPNs in the PD network.

We traced the directed cliques up to dimension 13 in our model. In the healthy PD0 model 6-dimensional cliques were the most abundant (10^7^) whereas 5-dimensional cliques were most numerous (10^5^) at the PD2 stage. In disease states (especially in PD2 and PD3) the number of directed cliques and their dimension drastically decreased (see Figure 4B) because of SPN dendritic atrophy. However cliques of dimension up to 13 are present in PD1 while the maximal clique dimension we found in PD0 is 12 (see Figure 4B). The mechanism underlying the formation of new higher dimensional cliques in PD1 is for example as follows: consider a 2-clique in PD0 consisting of two iSPNs and one dSPN. Now if, in PD1, a sprouted FS axon projects to all neurons in this clique it will create a 3-clique with FS as the source (schematized in Figure 4C). Just as in this example, because of their connection probability, FS are in the perfect position to form cliques of higher dimensions (not containing ChIN). In particular FS are not postsynaptic to any other neuron type in the network (Figure 1C), so when they belong to a clique, one FS is always the source. In summary, SPN dendritic atrophy and FS axonal growth counter each other and only in PD1 condition we observed an increase in maximum clique dimension (while the total number of cliques in PD1 was lower that in PD0).

Interneurons, despite constituting only 5% of the striatal neuron population, have a key role in maintaining higher order network connectivity, especially during PD progression, as shown in this study. To illustrate their role in directed clique formation we ablated all different types of interneurons (FS, LTS and ChIN) from the network. In the healthy network, ablation of interneurons did not drastically affect the count distribution of cliques (Figure 4B in log-scale, black solid and dotted line). However, removal of all types of interneurons drastically reduced both the count as well as the maximum clique dimension in PD networks (Figure 4B dotted lines).

If the increase in the FS-iSPN connectivity reported in Gittis et al. (2011) was not modelled (following the results in Gomez et al. (2019); Salin et al. (2009)), new cliques having FS as source were not formed and there was no increase in the maximal clique dimension in PD1 compared to PD0. Nevertheless, even in this case the interneurons ablation substantially reduced both clique count and clique dimension in PD networks.

The specific effect that SPN dendritic atrophy have on the directed clique distribution was further investigated by comparing the healthy PD0 and PD networks not including interneurons (and hence containing only SPNs, dashed lines in Figure 4B). Loss of distal dendrites during the PD stages resulted in a decrease in the number of SPN-SPN synapses. To explore the relation between dendritic loss and synapses reduction, we randomly removed synapses from the healthy (interneuron ablated) PD0 network to match the number of those (synapses) left in the PD stages and compared the distribution of the directed clique (Supplementary Figure 3). The removal of both proximal and distal synapses during the random synaptic erosion caused a more dramatic decrease in both the clique count and clique dimension compared to the distribution of directed cliques when mainly distal synapses were lost (because located on the degenerated dendrites). This indicates that proximal synapses contribute more to clique formation than distal ones.

#### Composition of directed cliques

To better understand which types of cliques are affected during PD progression we categorised cliques by their composition type as: all dSPNs, all iSPNs, at least one interneuron, and dSPN and iSPN (for comparison between PD0 and PD2 see Figure 5 A; for comparison between all the PD stages in dimension 3 and 5 see Figure 5 B, C and Supplementary Figure 4 for cliques of other dimensions). Most directed cliques in the healthy network PD0 were exclusively formed by iSPNs and reached dimension 12 while in PD2 the majority of cliques contained at least one interneuron (FS, LTS, ChIN) and reached dimension 10 (see Figure 5A). Without interneurons in the PD2 network the maximum clique dimension was only 6. These results show that during PD progression, cliques without interneurons were clearly more affected and decreased at a faster rate. In fact, cliques of dimension 5 either only containing dSPNs or dSPNs and iSPNs were absent in PD3 (Figure 5 C). From dimension 3, the cliques containing at least one interneuron are always the most abundant in PD (See supplementary Figure 4).

**Figure 5.**
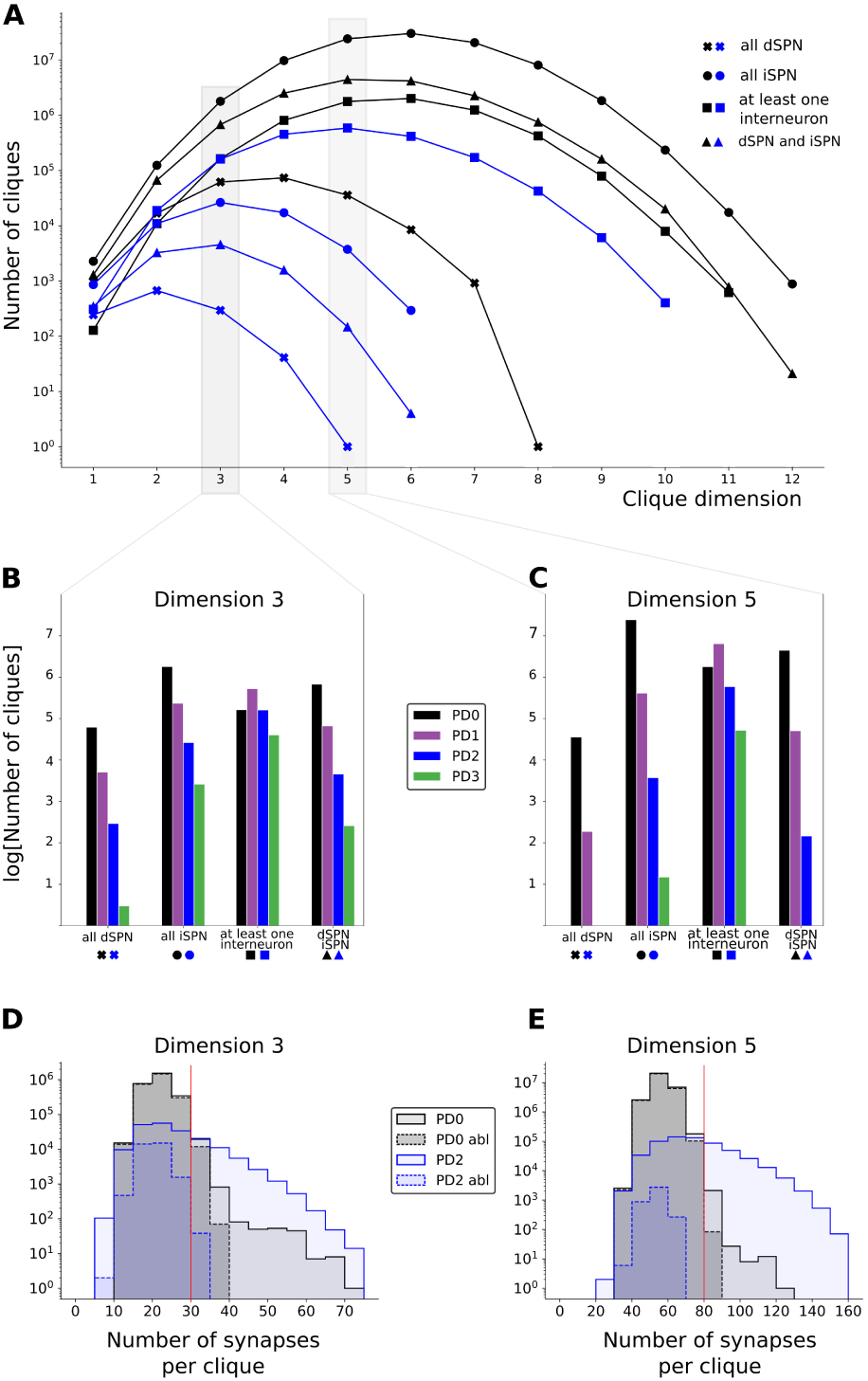
Composition of directed cliques. (A, B and C) The presence of directed cliques composed by only dSPN cells (x marker), only iSPN cells (circle marker), containing at least one interneuron (square marker), containing both dSPN and iSPN (triangular marker) is analysed. (A) Number, in log scale, of directed cliques with specific neuron compo-sitions described above as a function of the clique dimension in the healthy network PD0 (black curves) and at Parkinsonian stage PD2 (blue curves). (B,C) Number, in log scale, of cliques in dimension 3 and dimension 5, respectively in PD0 (in black), PD1 (in purple), PD2 (in blue) and PD3 (in green) subdivided within the specific neuron compositions. (D,E) represent the log scale histogram of cliques in dimension 3 and 5 with a given number of synapses, respectively, in PD0 (light grey), PD0 interneuron ablated (dark grey dashed boundary), PD2 (light blue) and PD2 interneuron ablated (light blue dashed boundary). Vertical red lines represent the thresholds such that cliques with a subthreshold number of synapses were more abundant in PD0, while cliques with a suprathreshold number of synapses were more abundant in PD2.

Directed cliques were further characterised by the number of synapses between all pairs of neurons composing the cliques (Figure 5 D, E). We found a “threshold” (30 for cliques in dimension 3 and 80 for cliques in dimension 5) such that cliques with a subthreshold number of synapses were more abundant in PD0, while cliques with a suprathreshold number of synapses, although generally fewer, were more abundant in PD2 (Figure 5 D, E: notice that below the threshold the black curve representing PD0 is above the blue curve representing PD2 and above the threshold the opposite holds). Moreover in the ablated networks the number of synapses per clique decreases faster in PD2 than in PD0 (Figure 5 D, E).

#### The role of interneurons in network high connectivity

To confirm that interneurons are crucial for maintaining the dimensionality of the cliques we progressively and randomly pruned the PD networks in two different ways and compared the results to that obtained in the corresponding ablated networks (i.e. the network without interneurons already represented in Figure 4B). First, we pruned synapses from the entire network (including every cell type). Second, we only pruned SPN-SPN synapses (Figure 6 A, B, C and D, E, F, respectively). Because of the interneurons’ involvement in high dimensional cliques, if their connectivity is kept fixed and only the SPN connectivity is eroded (as in the second erosion) the maximal clique dimension is expected to be greater than the maximal of the interneuron ablated networks.

**Figure 6.**
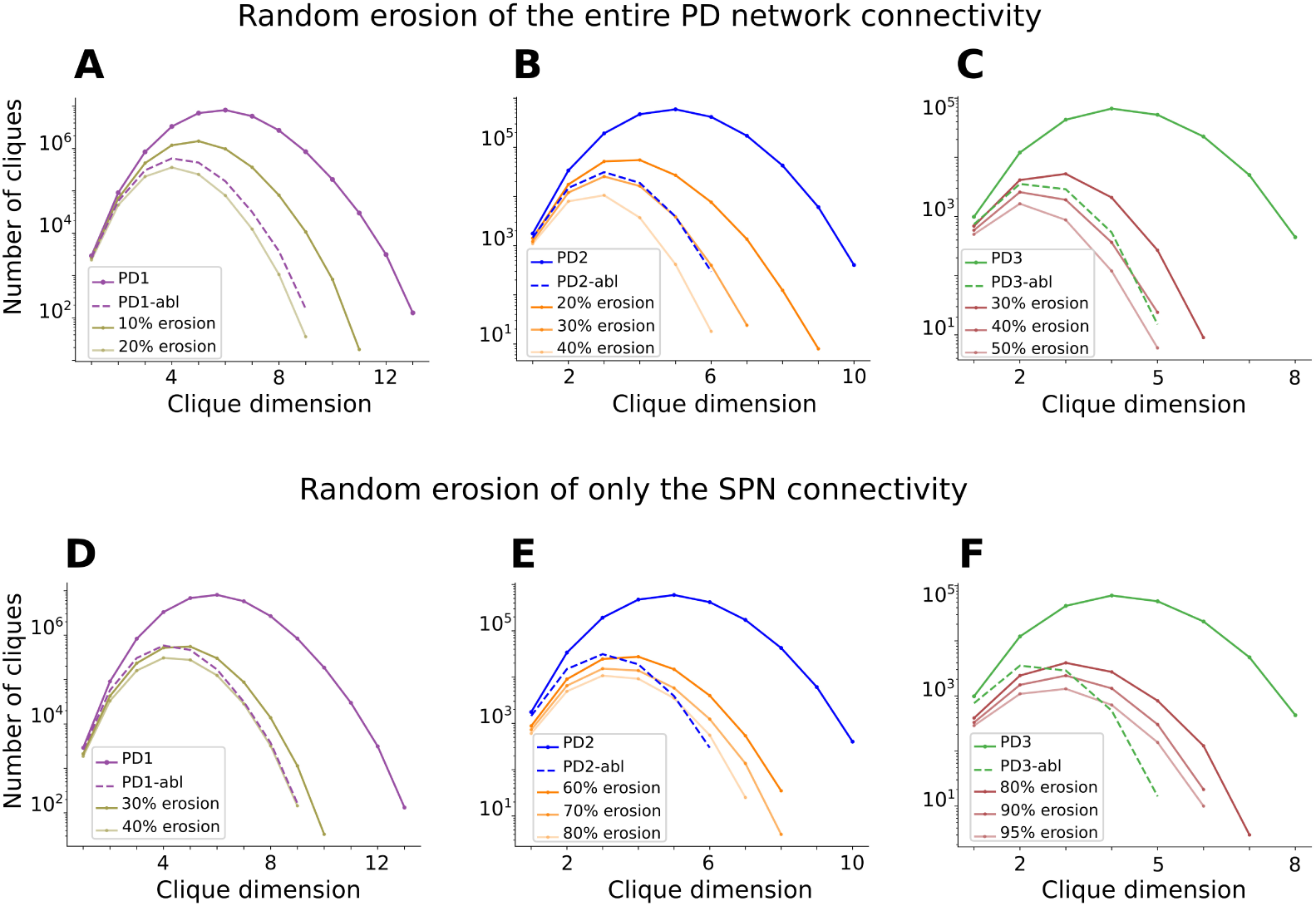
Interneurons are key to maintaining network connectivity. Despite being only 5% of the neuron population, ablation of the interneurons leads to significant loss of connectivity in each PD stage. The importance of the interneurons can be observed by assessing how many random synapses (directed edges) in the network have to be removed to obtain a comparable effect to ablating the interneurons on directed clique counts. In PD1 (A), 10-20% of connections need to be eroded; in PD2 (B) around 30%, while in PD3 (C) approximately 40%. If instead only SPN synapses are removed, the fraction of synapses that need to be removed is even higher. In PD1 (D), 30-40% of the SPNs synapses need to be eroded, in PD2 (E) between 60-80% and in PD3 (F) 80-95%.

When starting from PD1, to mirror the directed clique count of the interneuron ablated (PD1) network, between 10-20% of the synapses had to be removed when eroding the entire network (Figure 6A) while between 30-40% of the connections had to be eroded when removing only the SPN connectivity (Figure 6D). These percentages are expected to increase when considering more severe PD stages. In PD2 and PD3 around 30% and 40% erosion of the network respectively (Figure 6B-C) was needed to mimic the corresponding interneuron ablation. As expected, even when eroding only the SPN connectivity by 80% (PD2, Figure 6E) and 95% (PD3, Figure 6F), the maximal clique dimension obtained was one dimension greater than the corresponding maximal dimension in the interneuron ablated networks.

Independently on the synaptic erosion setting, it was possible to match the maximal number of cliques of the interneuron ablated networks with an eroded network. However, because of the discrepancy in the maximal clique dimensions, with clique dimensions dropping in the interneuron ablated networks, the shapes of the clique distributions of the eroded networks only match when all synapse types are included in the erosion process.

### Transfer of cortical input to striatal output

Another direct consequence of dendritic atrophy is loss of glutamatergic inputs. To investigate how this may affect the transfer of input from cortex (and thalamus), we simulated a set of Parkinsonian dSPN and iSPN and compared the neuron firing rate for different types of inputs with the healthy counterparts.

To this end, we used dSPN and iSPN which were tuned to reproduce physiological changes measured in Fieblinger et al. (2014) (see Materials and Methods). The dSPN PD2 electrophysiological models accounted for the increase in dSPN intrinsic excitability (Figure 7 top) while the iSPN PD2 models accounted for the decrease in iSPN intrinsic excitability (Figure 7 bottom).

**Figure 7.**
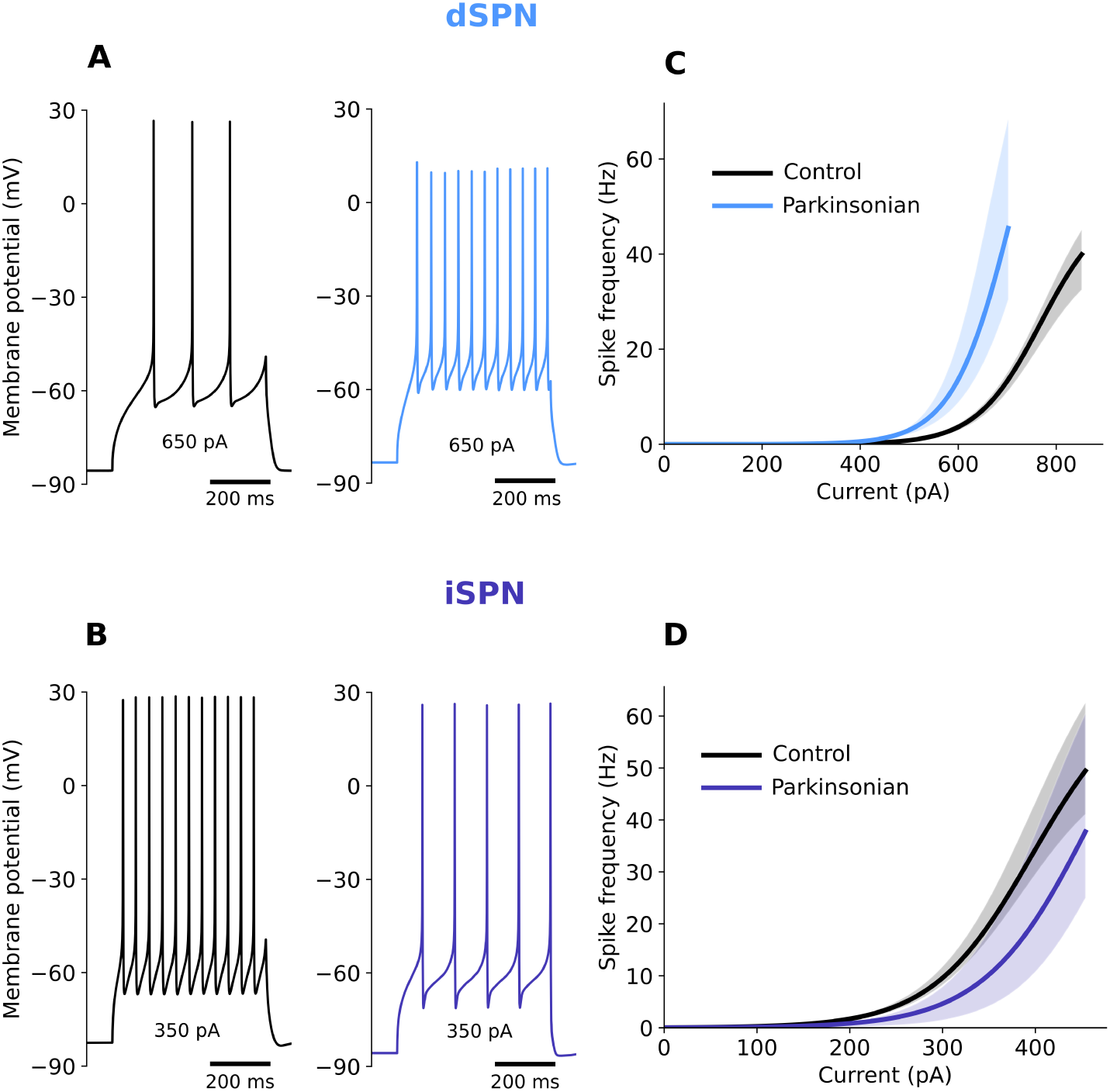
Modelling of the electrophysiological properties of SPNs during PD. Changes of excitability and shape of the action potential in the striatal projection neurons’ model of the direct pathway (dSPN, A) and indirect pathway (iSPN, B). Voltage traces and current-frequency response curves are shown for healthy neurons (Control, black lines) and neuron models adjusted to mimic physiological changes typical for PD (Parkinsonian, colour lines). Voltage plots illustrate discharge patterns of the healthy and PD cells in response to the same somatic direct current injection. Current-frequency curves are shown for the single-cell models, dSPN (C) and iSPN (D), using one morphology (up to 9 variations) for each cell type and multiple fitted electrical parameter sets (up to 10 for each cell). Shaded regions represent range values.

These models were then used to investigate the transfer of cortico-striatal inputs to the striatum. The number of cortico-striatal synapses (=size of the input ensemble) in the control case (PD0) was tuned for each morphology to obtain an output frequency of about 10 Hz when the synapses were receiving 5 Hz of Poisson type spiking input (see Materials and Methods and Supplementary Figure 5). For the PD2 network we used *Snudda* to estimate the number of synapses remaining on the neurons after dendritic atrophy.

The activation of the (remaining) synapses on the PD2 morphologies (Figure 8A, red circles) were not sufficient to equalise the output frequency obtained in healthy SPNs for the same synaptic input frequency. Neurons in the PD2 stage spiked at a very low firing rate (Figure 8B-E compare grey and blue lines). Therefore, we used two strategies to “compensate” for the loss of synapses and restore the activity level to the PD0 case: the remaining synapses were strengthened by increasing their conductance (Figure 8A, middle panel) or the synapses on the atrophied dendrites were recovered and distributed over the remaining dendrites (synapses rewiring; Figure 8A, right panel). These two forms of compensations were done gradually to better quantify their effect (see Materials and Methods) and a schematization of the settings is illustrated in Figure 8A.

**Figure 8.**
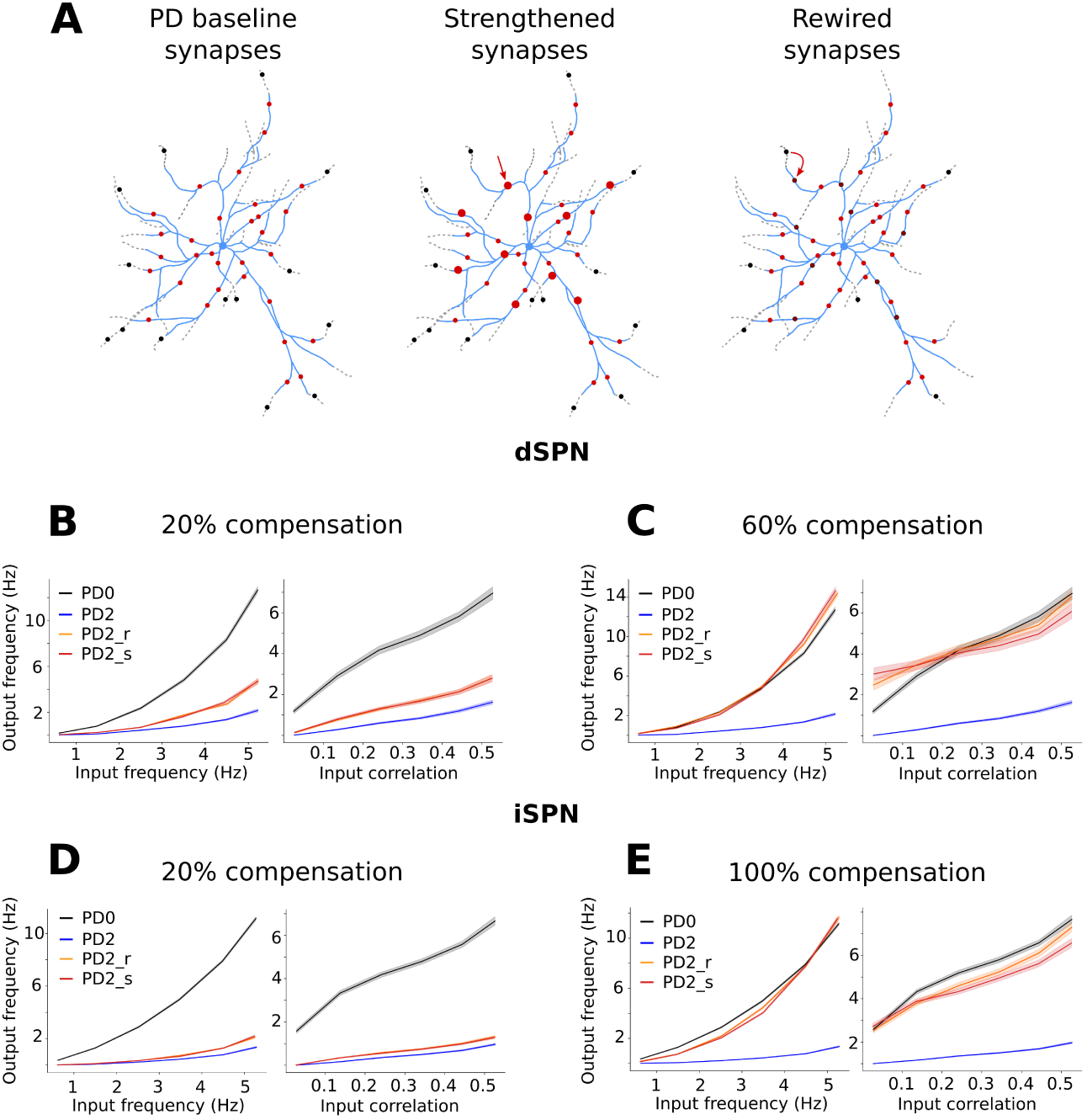
Striatal projection neurons (SPNs) responses to stimulation of cortico-striatal synapses in the control network (PD0) and during Parkin-son’s disease (PD2). (A) Because of the dendritic atrophy of SPNs (dotted grey branches), some cortico-striatal synapses are lost (black circles). The synapses (red circles) on the remaining branches (light blue) are not sufficient to produce the same response as the healthy cells. For this reason two compensatory mechanisms were implemented to restore the activity level. Percentages of the remaining synapses (20%, 40%, 60%, 80% and 100%) were strengthened (illustrated as larger circles in middle panel) or percentages of lost synapses were rewired (recovered and redistributed) on the remaining dendrites (illustrated as dark red circles in right panel). Different synaptic input frequencies (from 0.5 Hz to 5 Hz in steps of 0.5 Hz) and input correlations (from 0 to 0.5 in steps of 0.1) were used to stimulate the neurons. Specifically all the combinations were considered and the output frequency is plotted either as function of the input frequency averaging over the input correlations (left panels B, C, D, E) or as function of the input correlations averaging over the input frequencies (right panels B, C, D, E). Line colours represent the output frequency of the healthy (PD0, black) neurons, the Parkinsonian ones (PD2, blue) as well as levels of rewiring (PD2 r, orange) and strengthening (PD2 s, red). Specifically, (B) and (C) refer to dSPN when 20% and 60% compensation were applied respectively. (D) and (E) refer to iSPN when 20% and 100% compensation were applied respectively.

With this setup we systematically varied the SPN glutamatergic synaptic activation frequency as well as their correlation (60 different input configurations) and measured the output firing rate of all the SPNs in their healthy state (PD0) and unhealthy states (PD2) with 10 types of compensations (see Materials and Methods for details). Either rewiring or strengthening of synapses gave quite similar results, and was sufficient to match the output average firing rate of the PD2 neurons with that of their healthy counterparts. Notably, given the difference in SPN excitability (Figure 7), smaller compensation was needed for dSPNs as compared to the iSPNs (compare Figure 8C, E). Typically, increase in correlations results in higher output firing rate. However, for dSPNs both during synaptic rewiring and strengthening, when the compensation accounted for 60% or more, increased correlation resulted in a reduction in the input firing rate (Supplementary Figure 5 and Supplementary Figure 8 for a conceptual explanation).

In summary, these results highlight that strengthening and rewiring (with and without correlations) have quite similar effects here, and that especially iSPNs would need more compensation to approximate the output frequency obtained in the healthy case. If taking into account that in PD synaptic density may be decreased also on the remaining dendrites Villalba and Smith (2018), the enhanced glutamatergic effects are predicted to be even larger. However, here we have only focused on the feedforward transfer of cortical inputs to SPNs. Results may change if we consider both disease related changes in feedforward (due to FS axon sprouting) and recurrent inhibition (due to SPN dendrite atrophy).

## DISCUSSION

Parkinson’s disease leads to several changes in the striatal neurons, morphologically and electrophysiologically. We investigated how the effects of PD on striatal neurons impact the richness of the striatal network connectivity and also predicted resulting increases of the effective glutamatergic drive (in accordance with glutamatergic hyperactivity Campanelli, Natale, Marino, Ghiglieri, and Calabresi (2022)). The innovation of this study lies in the use of an *in silico* biophysically detailed striatal microcircuit model to disentangle the morphological and electrophysiological effects, something that currently would be a challenging task to asses in an experimental setting.

Progression of PD is accompanied by gradual changes in the neuron morphologies and intrastriatal synaptic connections. To understand how these changes affect the connectivity structure of the local striatal network we focused on directed cliques, a tool from computational topology. Here we observed that the number and dimension of directed cliques decreased as the disease progressed, highlighting the degeneration of higher order network connectivity structure. Unexpectedly, we also found that interneurons are crucial in both maintaining the network connectivity during PD and in the formation of high dimensional cliques (in particular due to sprouting of the FS axons). Most of the high dimensional cliques in the PD networks (dimension 3 or higher in PD1, dimension 2 or higher in PD2 and PD3 stages) contained at least one interneuron. Our clique count curves have shapes similar to those reported for the neocortical networks (Reimann et al. (2017) and Kanari et al. (2022)). However, there are notable quantitative differences between a striatal and a neocortical network in the number of cliques, maximal dimensions and behaviour of the curves when considering networks of degenerated neurons. These differences are likely to be due to the differences in the neuron types, pairwise connectivity, spatial distribution of neurons (e.g. layering in the neocortex) and strategies for modelling neuronal degeneration. Our PD morphologies are directly degenerated (or grown) using the software *treem*, while other methods such as that used in Kanari et al. (2022) base their degeneration on topological descriptors of the dendritic tree. Directed cliques indicate convergence of activity and therefore their presence in a network could imply emergence of synchrony in the network. Indeed, consistent with this, Reimann et al. (2017) related directed cliques in a neocortical microcircuit model to synchrony following stimulus from thalamus. However, the exact relationship between clique count and degree of correlations is not well understood. Moreover, in the striatum recurrent connections are GABAergic (inhibitory) and it is unclear whether directed cliques with inhibitory synapses have the same impact on network activity as the directed cliques with mainly excitatory synapses. While a full simulation of striatal activity dynamics in different stages of PD is beyond the scope of the current initial work, our microcircuit model framework can be used to investigate the effects of directed clique formation on striatal activity in future studies. In this scenario, the altered properties of ChIN and LTS interneurons can be included (Shen, Zhai, and Surmeier (2022)).

Chronic dopamine depletion also induces many cell specific alterations in intrinsic excitability and glutamatergic synaptic connectivity. It is worth mentioning that glutamate signaling contributes to the degeneration. Indeed, decortication (in rodents models) counteracts the morphological alterations (Zhu et al. (2019)). In healthy conditions a transient dopamine increase enhances the excitability of dSPN via D1 dopamine receptor signalling, while it decreases the excitability of iSPN through D2 dopamine receptor signalling (Surmeier, Graves, and Shen (2014)). Dopamine depletion in PD leads to increased intrinsic excitability in dSPNs (assessed via somatic current injection) and decreased excitability in iSPNs (Fieblinger et al. (2014), Ketzef et al. (2017)). Higher neuron excitability does not automatically imply higher firing rates *in vivo*, as network interactions, neuromodulation and various synaptic alterations can affect the final firing rate. How firing rates of dSPNs and iSPNs are altered *in vivo* during PD in response to ongoing cortical (and thalamic) activity is not clear from the literature. Conflicting results for dSPN and iSPN firing rates have been reported both in PD animal models and human patients (Singh et al. (2016), Beck, Singh, and Papa (2018), Valsky et al. (2020)). Several labs have reported that iSPNs have higher firing rate than dSPN in both anaesthetised and awake rats (Chen, Morales, Woodward, Hoffer, and Janak (2001), Mallet, Ballion, Le Moine, and Gonon (2006), Sharott et al. (2017)). Similarly, Parker et al. (2018) have reported higher Ca^2+^ event frequency in iSPNs compared to dSPNs during spontaneous activity in a mouse model of PD. However, Maltese, March, Bashaw, and Tritsch (2021) reported no significant difference between dSPN and iSPN Ca^2+^ event frequency. Consistent with this, the spike firing rate of dSPNs and iSPNs in DA-depleted mice was comparable in both anaesthetised (Ketzef et al. (2017)) and awake (de la Torre-Martinez et al. (2023)) states. Importantly for our study is that, to the best of our knowledge, no one has reported decreased activity in any of the SPN neuron types, although the ensemble size of dSPNs might be decreased *in vivo* (in mice) (Maltese et al. (2021)). In our *in-silico* simulations of SPN glutamatergic activation we accounted for both the SPN morphological degenerations and intrinsic excitability changes. We found that Parkinsonian SPNs firing rates were much lower than their healthy counterparts while keeping the synaptic densities on the non-degenerated (remaining) dendritic branches and the individual synaptic strengths unchanged. This is a consequence of the majority of the spines being located on distal dendrites which account for the most part of the total dendritic length (Hjorth et al. (2020)). To investigate how to restore the firing rate of SPNs to at least their healthy level, we tested two different scenarios: a) strengthening of the remaining synapses, and b) rewiring the lost synapses from the degenerated dendritic branches onto the surviving dendrites. For both these scenarios, we studied how the SPNs spike rate is influenced by several different combinations of synaptic input correlations and input frequencies. Interestingly, dSPNs and iSPNs needed different levels of compensation, in particular iSPNs needed more or stronger glutamatergic drive (60% and 100% recovery, respectively). Thus, our result predicts that the glutamatergic drive must undergo large quantitative changes during chronic dopamine depletion. This is in line with the glutamatergic hyperactivity (Campanelli et al. (2022)). Several experimental studies have reported mechanisms that might contribute to a functionally more effective glutamatergic drive during dopamine depletion. While there is not much support for increased averaged spiking activity in cortex, rather the opposite (Underwood and Parr-Brownlie (2021), Viaro, Morari, and Franchi (2011), Bamford et al. (2004)), bursting develops and this might activate striatal neurons more effectively (Cagnan et al. (2019)). Furthermore, the parafascicular thalamic nucleus to iSPN drive is increased (Tanimura, Du, Kondapalli, Wokosin, and Surmeier (2019)), while at the same time, the intrastriatal lateral inhibition is decreased (López-Huerta et al. (2013), Taverna et al. (2008)). In addition, while dopamine depletion itself might decrease presynaptic inhibition (Bamford et al. (2004)), it also leads to an increase in the striatal acetylcholine levels (Ztaou and Amalric (2019), Aosaki, Miura, Suzuki, Nishimura, and Masuda (2010), Ding et al. (2006)) (that might depolarise the SPNs via M1 cholinergic receptors *in vivo*), and alterations in the dynamics of the burst-pause response in ChINs (as this partly depends on D2R and D5R receptor activation). The latter might perhaps decrease presynaptic cortico-striatal inhibition (Aosaki et al. (2010), Pancani et al. (2014)). Moreover, the downregulation of glutamate transporters in striatal glia cells (Chung, Chen, Chan, and Yung (2008)), the enhancement of some NMDA subunit types in the membrane compartment (Gan, Qi, Mao, and Liu (2014)), and a change in the SPN A-type K^+^ ion channel conductance and dynamics could together produce larger summation of EPSPs in the SPN dendrites (Azdad et al. (2009)). Given our prediction of the significant amount of extra glutamatergic drive needed (60-100%) to at least allow the parkinsonian SPNs to fire as much as in the healthy state, it will be important to better quantify experimentally how these types of observed alterations contribute to the enhanced activation of striatal SPNs.

Using an *in-silico* microcircuitry gives the advantage of making clear modelling assumptions and testing different scenarios to generate predictions as well as new questions. Moreover, using *in-silico* reconstructions allowed us to disentangle the effect of the morphological and resulting network topological alterations from the more complex electrophysiological changes that the different neuron types undergo. Our Python code is open source with reproducible workflows that others can explore with modified assumptions.

We made predictions on how the directed clique count changes during PD, and although challenging, these could be measured experimentally. We also predicted that chronic dopamine depletion in PD significantly increases the effective glutamatergic drive, especially to iSPNs. The glutamatergic hyperactivity is one significant driver of several of the morphological changes seen (Zhu et al. (2019)). Targeting some of the most contributing factors may be relevant for counteracting PD progression. For example, preventing or reversing glutamatergic hyperactivity might prevent alterations in the SPN morphology as well as the local network topology. Future experiments might shed light on which of the PD progression mechanisms has the largest impact on symptoms and whether mechanism-specific treatments at certain stages of PD could slow down the progression of the disease.

In summary, our work highlighted that just measuring the pairwise connectivity between neurons gives an incomplete description of the network connectivity. Here we did not assume neurons to connect in a completely random fashion, instead we used the neuron morphologies to further constrain and predict the connectivity. We showed that directed cliques provided a richer characterization of the predicted changes in the network structure with respect to PD progression. We highlighted that the glutamatergic drive on the remaining synapses onto SPNs must undergo large quantitative increases to compensate for the effects of chronic dopamine depletion. Moreover the extent of these alterations should be quite different between dSPN and iSPN.

## MATERIALS AND METHODS

Box and arrow diagrams in Supplementary Figure 9 summarise this section, highlighting the software, tools and formalisms used.

### Network creation

To investigate the striatal circuitry, 100 000 neurons were placed in a defined volume with appropriate cell densities (approximately 80 500 neurons/mm^3^) using the simulation environment *Snudda*, which allows to predict synaptic connectivity based on touch detection and a set of pruning rules (for details see Hjorth et al. (2020) and Hjorth et al. (2021)). The touch-detection process located putative synapses based on close proximity between axons and dendrites/soma of different neurons. During pruning, the number of synapses between pairs was reduced based on rules implemented to match pairwise connectivity data (number of synapses and connection probability between neuron types). The pruning rules, inspired by Reimann et al. (2015), were the following: a random fraction of all synapses was removed; synapses were removed according to distance-dependent functions (to match FS and LTS interneurons preference to make proximal and distal synapses on SPNs, respectively); if an excessive number of synapses between neuron pairs was detected, a fraction of them was removed; if too few synapses were detected between neuron pairs, they were all removed with a certain probability (that increased the fewer synapses connected the neurons); all the synapses between coupled neurons were removed based on a specified probability.

Here the goal is to create both a healthy wild type (PD0) network and a network representing the progression of Parkinson disease (PD1, PD2 and PD3).

### Converting a healthy network into a Parkinsonian network

#### Change in neuron morphologies

During Parkinson’s disease the striatal projection neurons’ (SPN) dendrites degenerate, causing a reduction in the number of distal synapses. Fast-spiking (FS) axons, in turn, grow leading to an addition of GABAergic synapses. These two effects contribute to changing the connectivity of the network. Neurodegeneration was modelled as a progressive loss of the most distal fragments of the dendritic arbors of the SPN. This process resulted in systematic decrease of the total dendritic length while not much affecting the maximum radius of the dendritic field and the number of primary dendrites similar to the data in Fieblinger et al. (2014). The latter reports reduction of the total dendritic length to 75.8% and 69.7% in dSPN and iSPN populations, respectively. Number of primary dendrites reduced significantly in dSPNs from 8 to 6 and less noticeably in iSPNs from 6 to 5, which was within the range of control values. Similarly, the number of the dendritic branching points dropped to 69.4% in dSPNs and was unchanged in iSPNs. In our study, morphological reconstructions were manipulated using the Python module *treem* (Kozlov, A. K., 2021, Hjorth et al. (2021)). The initial PD0 morphologies were labelled PD0. Dendritic arbors were sampled at a fixed spatial resolution of 3 µm. SPN dendrites were shortened step-wise to mimic degeneration so that at each step of the algorithm one dendritic segment 3 µm long is truncated at every terminal. Morphologies after 10 and 20 truncation steps were labelled PD1 and PD2, respectively. The PD1 and PD2 neuron degenerations were based on mouse data from Fieblinger et al. (2014), while PD3 (30 steps) corresponds to a greater dendritic loss resembling what the human cells exhibit. Changes in the mean total dendritic length in our model are outlined in Table 1 and illustrated in Figure 2A-C. To mimic the rapid growth of FS axons, we extended each axonal terminal of PD0 FS morphologies by 61 µm and kept them unchanged between pathological PD stages 1-3 (See Table 1 and (Figure 2D-E).

**Table 1.** Changes in the mean total dendritic and axonal length in the model. Striatal projection neurons (dSPN and iSPN) undergo dendritic degeneration during Parkinson’s disease (PD1, PD2 and PD3). fast spiking interneurons (FS) may instead exhibit axonal sprouting.

| Neuron type | PD0 | PD1 | PD2 | PD3 |
| --- | --- | --- | --- | --- |
| Mean total dendritic length |  |  |  |  |
| dSPN | 3997.9 $\mu\text{m}$ (100%) | 2890.8 $\mu\text{m}$ (72%) | 1984.6 $\mu\text{m}$ (50%) | 1288.6 $\mu\text{m}$ (32%) |
| iSPN | 3116.7 $\mu\text{m}$ (100%) | 2362.5 $\mu\text{m}$ (76%) | 1774.5 $\mu\text{m}$ (57%) | 1302 $\mu\text{m}$ (42%) |
| Mean total axonal length |  |  |  |  |
| FS | 25572 $\mu\text{m}$ (100%) | 41249 $\mu\text{m}$ (161.3%) | | |

#### Evolution of the striatal network structure during PD

There are different ways to generate the PD network. One approach is to start from the complete PD0 network, and remove the synapses that were placed on dendritic branches lost during the degeneration of the morphologies (degeneration method). This was done by swapping the PD0 morphologies for their corresponding PD morphologies, keeping the location and orientation and identifying which synapses are no longer attached to a dendrite. Another approach is to start with the PD morphologies in the corresponding locations and perform a new touch detection to determine where the synapses are (de novo method). In the first case the degenerated synapses were in the same location as before, but the extra FS synapses were missing. In the second case, the extra FS synapses were included, but the remaining synapses were not in the same location as before. To compensate for this, a hybrid method was implemented, where the synapses from the degeneration method and the de novo method were combined. In addition, there was also a difference in the number of synapses detected between the two methods, since in the first case pruning was done before degeneration, and in the second case degeneration was done before pruning, (see Supplementary Figure 1). In the hybrid method the number of synapses between neuron types was tuned to match those detected in the de novo method. The fraction of synapses remapped, i.e. synapses taken from the de novo method and added to the degenerated method’s set of synapses, is called the remapping fraction. In summary, the hybrid method retained the position for the remaining synapses, while adding the new FS synapses to the network. It also retained a comparable number of synapses as a de novo detected PD network.

### Topological measurements

Directed cliques can be used to measure higher order connectivity patterns in directed graphs. Following Reimann et al. (2017) a directed graph is defined as a pair of sets (*V, E*) equipped with an injective function *τ* : *E → V × V* where *V* is the set of vertices, *E* is the set of directed edges and assigns to a directed edge its source and target respectively. Furthermore it is assumed that a directed graph has no self loops (i.e. if *τ* (*e*) = (*v*_1_*, v*_2_) then *v*_1_ *̸*= *v*_2_). If *τ* (*e*) = (*v*_1_*, v*_2_), we call *e* a directed edge from *v*_1_ to *v*_2_. Notice that the injectivity assumption implies that it is not possible to have more than one directed edge from *v*_1_ to *v*_2_. Two vertices *v*_1_ and *v*_2_ can however be reciprocally connected with one directed edge from *v*_1_ to *v*_2_ and one directed edge from *v*_2_ to *v*_1_. Two vertices *v*_1_ and *v*_2_ in a directed graph are said to be connected if there is either an edge from *v*_1_ to *v*_2_, or an edge from *v*_2_ to *v*_1_, or both. A vertex *v* of a directed graph is called a *source* if it can only be the source of one or more directed edges, i.e. *v* can only appear in the first coordinate of the image of the function. In the opposite way, a vertex *w* is a *sink* if it can only be the target of one or more directed edges, i.e. *w* can only appear in the second coordinate of the image of the function. Informally one can say that all the edges containing a source are from this vertex and all the edges containing a sink are to this vertex. A directed clique is a set of vertices in a directed graph which are all to all connected and there exists a unique source and a unique sink among them. A directed clique consisting of *n* + 1 vertices is called a clique of dimension *n* or directed *n*-clique. If a partial order is defined on the vertices of a directed graph, where *v*_1_ *< v*_2_ if there is an edge from *v*_1_ to *v*_2_, a directed clique is a totally ordered subset of the vertices whose smallest element is the source and the largest element the sink. Directed cliques can be thought of as feedforward motifs from the source to the sink (See Figure 4 A1-4). In this article a directed graph is a structural network of neurons where vertices represent neurons and a directed edge represents the presence of synapses connecting a presynaptic neuron to a postsynaptic neuron. The source of a directed clique can then be seen as a neuron which is presynaptic to all the other neurons in the clique while the sink is a neuron which is postsynaptic to all the other neurons in the clique.

### Selection of the core of the network

To avoid underestimating the connections and consequently the number and or types of cliques, it is important for the analysis that the neurons investigated have all their connected pre- and post-synaptic partners included in the network. In other words, to prevent potential edge effects a set of neurons (referred to as *kernel* neurons here) at the centre of the network is selected. All the neurons (both pre- and post-synaptic) connected to the kernel neurons were identified and labelled as the *core* together with the kernel neurons. Without this the connectivity for the neurons included in the analysis would be underestimated. Directed clique analysis was performed for neurons in the core and all cliques were required to have at least one neuron belonging to the set of kernel neurons. All the results shown were obtained using a kernel of 8 neurons, 4 dSPNs and 4 iSPNs which resulted in a core of 2712 neurons (out of 100 000). The maximal distance between neurons in the kernel and their connected pre-or-postsynaptic neurons was around 550 µm. Also cores formed from kernels with exclusively dSPNs or iSPNs have been analysed, but because SPNs are intermixed within the striatum and present in equal number, a mixed kernel was preferred.

### Simulation of cortical input to dSPN and iSPN neurons

#### Models of striatal projection neurons

Computational models of the healthy dSPN and iSPN cells were taken from the previous studies (Hjorth et al. (2020), Hjorth et al. (2021)). We refer the reader to them for details of the equations describing the time evolution of the membrane voltage. Several neuron models of each type (n=4) were fitted to experimental data (Hjorth et al. (2020), Figure 2 and Figure S3). Every model was characterised by a unique dendritic morphology, rheobase current and a current-frequency relation. Evolutionary parameter fitting algorithm (Van Geit et al. (2016)) provides multiple electrical parameter sets (up to 10) for each neuron model. To introduce more physiological variability within the neuron populations, the optimised parameter sets were combined with modified dendritic morphologies (9 variations of each reconstruction using scaling factors from 0.6 to 1.4 and random rotations of the dendritic branches at the branching points). All morpho-electric combinations were then simulated and validated against physiological features of the experimental populations as in Hjorth et al. (2020).

#### Change in SPNs electrophysiology during PD

Stage PD2 of the cell morphology modification was used to model the mouse PD network throughout. Electric parameters of the model SPN cells were manually adjusted to reproduce physiological changes observed experimentally in 6-OHDA lesioned mice (Fieblinger et al. (2014)). Specifically, in dSPNs, there was a reduction in both rheobase and action potential amplitude, accompanied by an increase in input resistance. while in iSPNs the rheobase and afterhyperpolarization (AHP) were increased. Therefore, in dSPN models, both the fast sodium current and the transient potassium currents were reduced to account for the decrease in the action potential amplitude and AHP, the increase in excitability was mainly explained by the shorter total dendritic length (leading to higher input resistance of the neuron). Reduced excitability of iSPN cells in PD was achieved through strengthening of the inward rectifying potassium current and a corresponding increase of the leak conductance to maintain the unchanged resting membrane potential. Transient potassium current was also increased in PD iSPN’s to match the larger AHP. Voltage traces and current-frequency responses of the healthy and PD SPN models are shown in Figure 7.

### Setting and stimulating the input and compensatory mechanisms

Both healthy and Parkinsonian SPNs were simulated using the same cortico-striatal drive to investigate the change in their output frequency. The number of input spike trains (*n*) received by a neuron was determined by the neuron morphology and the type of compensation (see below). The number of synapses which were distributed on the PD0 morphologies was tuned to achieve an average output frequency of 10 Hz when the synapses were receiving 5 Hz of Poisson spike train without correlation. Using *Snudda* an estimate of the remaining synapses on the PD2 morphologies was obtained (47.5% and 53.25% for dSPN and iSPN respectively). Because of the reduction in the dendritic length and number of synapses (compared to PD0) the output frequencies of these neurons were lower than in the healthy counterparts. Hence, two compensatory strategies were implemented to restore the activity level to the PD0 case. Strengthening the (remaining) synapses by 20%, 40%, 60%, 80% and 100% and rewiring the lost synapses (which were located on the branches that atrophied) by the same percentages. The base synaptic conductance is 0.5 nS and during the strengthening first the difference between the total synaptic conductance in PD0 and PD2 was calculated (number of missing synapses times 0.5 nS), then a percentage of this value divided by the number of synapses in PD2 was added to the base conductance of the latter. For example, to strengthen the synapses by 20%, one has to compute:

▪ the total synaptic conductance in PD0 *→* cond_PD0_= n synPD0 *·* 0.5 nS,
▪ the total synaptic conductance in PD2 *→* cond_PD2_= n synPD2 *·* 0.5 nS,
▪ the difference *→* cond_PD0_ *−* cond_PD2_ = (n synPD0 *−* n synPD2) *·* 0.5 nS,
▪ add 20% of if to the total conductance in PD2 *→* str cond_PD2_ = cond_PD2_ + (cond_PD0_ *−* cond_PD2_) *·* 0.2.

This means that to strengthen the synapses in PD2 by x% the conductance of each synapse in PD2 has to be increased by 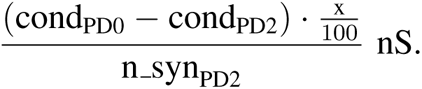 Hence in the 100% strengthening compensation step despite the number of synapses being different the total synaptic conductance is the same. Analogously, a percentage of the missing synapses was added at each step so that in the final case the number of synapses in PD2 was the same as PD0. Each input spike train was connected to the SPN with an excitatory synapse. Synapses were modelled using the Tsodyks*−*Markram formalisms (Markram, Wang, and Tsodyks (1998); Tsodyks and Markram (1997)) as in Hjorth et al. (2020). Synapses were activated using Poisson spike trains of average rate *λ* and pairwise correlation *c*. The pairwise correlation, *c*, of each input stream is generated by first creating a mother Poisson spike train of frequency, *f*, and n child Poisson spike trains of frequency (1 *− p*) *· f*. Where *p* = *c* is the probability that a mother spike is transferred to the child spike train. We systematically varied both *λ* [= 0.5 : 0.1 : 5 Hz] and *c* [= 0 : 0.1 : 0.5]. This resulted in 60 different combinations of input rate and correlations. All the 60 combinations between frequency and correlation were simulated for each stage (healthy, Parkinsonian, and 10 different compensations), for a total of 720 sets of simulations (per cell type). Each set consisted of several combinations of morphologies and electric parameters, and for each combination 10 simulations, which differed in the synapse distribution, were performed. The corresponding results are illustrated using heatmaps in Supplementary Figure 6 and 7 for dSPN and iSPN respectively. A single square in the heatmap represents the average over one set of simulations (which includes the different combinations of morphologies, parameters and synapse distributions). In summary, all the scenarios (PD0, PD2 and PD2 compensated) were simulated using all the possible combinations between frequency and correlation. For each pair (frequency, correlation) the stimulation lasted 10 seconds and a 2 second recovery period was included between pairs. In total 720 sets of simulations were reproduced (12 main scenarios including PD0, PD2, 5 strengthening and 5 rewiring times 10 different input frequencies times 6 different input correlations) for each combination of morphology and electric parameter. Each of this was repeated 10 times to include variability in the synaptic distributions.

### Simulation and data analysis tools

#### Treem

A neuron morphology processing software *Treem* (Kozlov (2021), https://github.com/a1eko/treem) provides data structure, programming interface and command-line tools for accessing and manipulating the digital reconstructions of the neuron morphology in Stockley-Wheal-Cannon format (SWC). Common operations that *treem* supports include *checking* the structural integrity of the reconstruction, *measuring* morphometric features, *repairing* cut branches, *correcting* for shrinkage, and *modifying* the morphology to model atrophy or growth.

We used this Python module, to modify the morphology to account for the changes that occur in PD i.e progressive degeneration of dendritic arbors in SPNs and growth of FS axons (see Material and Methods, Change in neuron morphologies).

#### Snudda

*Snudda* (https://github.com/Hjorthmedh/Snudda) is a modelling framework that enables the creation of large-scale networks by: placing neurons using appropriate density (within a user defined structure/volume), predicting synaptic connectivity based on touch detection and a set of pruning rules, setting up external input and neuromodulation, and finally running the simulations. It is written in Python and uses the NEURON simulator.

We used *Snudda* to create the networks (placement, touch detection and pruning) and setup the inputs. Neuron simulations were then performed using NEURON via *Snudda*.

#### Flagser

The number of directed cliques was computed using the software Flagser-count (https://github.com/JasonPSmith/flagser-count), an adaptation of Flagser (https://github.com/luetge/flagser) which deals with the more general problem of computing the homology and persistent homology of directed graphs. As a Python API for Flagser we refer to pyflagser (https://github.com/giotto-ai/pyflagser).

## ACKNOWLEDGMENTS

We thank: PhD Jason Smith for all the support regarding flagser-count; Tor Kjellsson Lindblom for the help setting up the simulations on Dardel; William Scott Thompson for the precious Blender advice; PhD Johanna Frost Nylén and PhD Yvonne Johansson for all the insightful conversations; and PhD Barbara Mahler for her support. Finally we thank Prof. Gilad Silberberg, Prof. Sten Grillner and Prof. Angela Cenci Nilsson for the enlightening discussions.

The computations were enabled by resources provided by the National Academic Infrastructure for Supercomputing in Sweden (NAISS) at PDC KTH partially funded by the Swedish Research Council through grant agreement no. 2022-06725.

## SUPPORTING INFORMATION

[pdf of Supplementary Material provided]

## REFERENCES

Alberquilla, S., Gonzalez-Granillo, A., Martín, E. D., & Moratalla, R. (2020). Dopamine regulates spine density in striatal projection neurons in a concentration-dependent manner. Neurobiology of disease, 134, 104666.

Aosaki, T., Miura, M., Suzuki, T., Nishimura, K., & Masuda, M. (2010). Acetylcholine–dopamine balance hypothesis in the striatum: An update. Geriatrics & gerontology international, 10, S148–S157.

Azdad, K., Chàvez, M., Bischop, P. D., Wetzelaer, P., Marescau, B., De Deyn, P. P., … Schiffmann, S. N. (2009). Homeostatic plasticity of striatal neurons intrinsic excitability following dopamine depletion. PloS one, 4(9), e6908.

Bamford, N. S., Zhang, H., Schmitz, Y., Wu, N.-P., Cepeda, C., Levine, M. S., … Sulzer, D. (2004). Heterosynaptic dopamine neurotransmission selects sets of corticostriatal terminals. Neuron, 42(4), 653–663.

Beck, G., Singh, A., & Papa, S. M. (2018). Dysregulation of striatal projection neurons in parkinson’s disease. Journal of Neural Transmission, 125, 449–460.

Bergman, H., Wichmann, T., Karmon, B., & DeLong, M. (1994). The primate subthalamic nucleus. ii. neuronal activity in the mptp model of parkinsonism. Journal of neurophysiology, 72(2), 507–520.

Burke, D. A., Rotstein, H. G., & Alvarez, V. A. (2017). Striatal local circuitry: a new framework for lateral inhibition. Neuron, 96(2), 267–284.

Cagnan, H., Mallet, N., Moll, C. K., Gulberti, A., Holt, A. B., Westphal, M., … others (2019). Temporal evolution of beta bursts in the parkinsonian cortical and basal ganglia network. Proceedings of the National Academy of Sciences, 116(32), 16095–16104.

Campanelli, F., Natale, G., Marino, G., Ghiglieri, V., & Calabresi, P. (2022). Striatal glutamatergic hyperactivity in parkinson’s disease. Neurobiology of Disease, 168, 105697.

Chen, M.-T., Morales, M., Woodward, D. J., Hoffer, B. J., & Janak, P. H. (2001). In vivo extracellular recording of striatal neurons in the awake rat following unilateral 6-hydroxydopamine lesions. Experimental neurology, 171(1), 72–83.

Chung, E., Chen, L., Chan, Y., & Yung, K. (2008). Downregulation of glial glutamate transporters after dopamine denervation in the striatum of 6-hydroxydopamine-lesioned rats. Journal of Comparative Neurology, 511(4), 421–437.

de la Torre-Martinez, R., Ketzef, M., & Silberberg, G. (2023). Ongoing movement controls sensory integration in the dorsolateral striatum. Nature Communications, 14(1), 1004.

Ding, J., Guzman, J. N., Tkatch, T., Chen, S., Goldberg, J. A., Ebert, P. J., … Surmeier, D. J. (2006). Rgs4-dependent attenuation of m4 autoreceptor function in striatal cholinergic interneurons following dopamine depletion. Nature neuroscience, 9(6), 832–842.

Doya, K. (2008). Modulators of decision making. Nature neuroscience, 11(4), 410–416.

Fieblinger, T., Graves, S. M., Sebel, L. E., Alcacer, C., Plotkin, J. L., Gertler, T. S., … others (2014). Cell type-specific plasticity of striatal projection neurons in parkinsonism and l-dopa-induced dyskinesia. Nature communications, 5(1), 5316.

Fieblinger, T., Zanetti, L., Sebastianutto, I., Breger, L., Quintino, L., Lockowandt, M., … Cenci, M. (2018). Striatonigral neurons divide into two distinct morphological-physiological phenotypes after chronic l-dopa treatment in parkinsonian rats. Scientific reports, 8(1), 10068.

Filipović, M., Ketzef, M., Reig, R., Aertsen, A., Silberberg, G., & Kumar, A. (2019). Direct pathway neurons in mouse dorsolateral striatum in vivo receive stronger synaptic input than indirect pathway neurons. Journal of neurophysiology, 122(6), 2294–2303.

Gan, J., Qi, C., Mao, L.-M., & Liu, Z. (2014). Changes in surface expression of n-methyl-d-aspartate receptors in the striatum in a rat model of parkinson’s disease. Drug design, development and therapy, 165–173.

Gittis, A. H., Hang, G. B., LaDow, E. S., Shoenfeld, L. R., Atallah, B. V., Finkbeiner, S., & Kreitzer, A. C. (2011). Rapid target-specific remodeling of fast-spiking inhibitory circuits after loss of dopamine. Neuron, 71(5), 858–868.

Gomez, G., Escande, M. V., Suarez, L., Rela, L., Belforte, J. E., Moratalla, R., … Taravini, I. R. E. (2019). Changes in dendritic spine density and inhibitory perisomatic connectivity onto medium spiny neurons in l-dopa-induced dyskinesia. Molecular Neurobiology, 56, 6261–6275.

Graveland, G., & DiFiglia, M. (1985). The frequency and distribution of medium–sized neurons with indented nuclei in the primate and rodent neostriatum. Brain Research, 327(1-2), 307–311.

Hjorth, J. J., Hellgren Kotaleski, J., & Kozlov, A. (2021). Predicting synaptic connectivity for large-scale microcircuit simulations using snudda. Neuroinformatics, 19(4), 685–701.

Hjorth, J. J., Kozlov, A., Carannante, I., Frost Nylén, J., Lindroos, R., Johansson, Y., … others (2020). The microcircuits of striatum in silico. Proceedings of the National Academy of Sciences, 117(17), 9554–9565.

Kanari, L., Dictus, H., Chalimourda, A., Arnaudon, A., Van Geit, W., Coste, B., … Markram, H. (2022). Computational synthesis of cortical dendritic morphologies. Cell Reports, 39(1).

Ketzef, M., Spigolon, G., Johansson, Y., Bonito-Oliva, A., Fisone, G., & Silberberg, G. (2017). Dopamine depletion impairs bilateral sensory processing in the striatum in a pathway-dependent manner. Neuron, 94(4), 855–865.

Kozlov, A. K. (2021). treem - neuron morphology processing tool. Zenodo. Retrieved from 10.5281/zenodo.4890844 doi: 10.5281/zenodo.4890844

López-Huerta, V. G., Carrillo-Reid, L., Galarraga, E., Tapia, D., Fiordelisio, T., Drucker-Colin, R., & Bargas, J. (2013). The balance of striatal feedback transmission is disrupted in a model of parkinsonism. Journal of Neuroscience, 33(11), 4964–4975.

Mallet, N., Ballion, B., Le Moine, C., & Gonon, F. (2006). Cortical inputs and gaba interneurons imbalance projection neurons in the striatum of parkinsonian rats. Journal of Neuroscience, 26(14), 3875–3884.

Mallet, N., Pogosyan, A., Márton, L. F., Bolam, J. P., Brown, P., & Magill, P. J. (2008). Parkinsonian beta oscillations in the external globus pallidus and their relationship with subthalamic nucleus activity. Journal of neuroscience, 28(52), 14245–14258.

Maltese, M., March, J. R., Bashaw, A. G., & Tritsch, N. X. (2021). Dopamine differentially modulates the size of projection neuron ensembles in the intact and dopamine-depleted striatum. Elife, 10, e68041.

Markram, H., Muller, E., Ramaswamy, S., Reimann, M. W., Abdellah, M., Sanchez, C. A., Schürmann, F. (2015). Reconstruction and simulation of neocortical microcircuitry. Cell, 163(2), 456–492.

Markram, H., Wang, Y., & Tsodyks, M. (1998). Differential signaling via the same axon of neocortical pyramidal neurons. Proceedings of the National Academy of Sciences, 95(9), 5323–5328.

McNeill, T. H., Brown, S. A., Rafols, J. A., & Shoulson, I. (1988). Atrophy of medium spiny i striatal dendrites in advanced parkinson’s disease. Brain research, 455(1), 148–152.

Pancani, T., Bolarinwa, C., Smith, Y., Lindsley, C. W., Conn, P. J., & Xiang, Z. (2014). M4 machr-mediated modulation of glutamatergic transmission at corticostriatal synapses. ACS chemical neuroscience, 5(4), 318–324.

Parker, J. G., Marshall, J. D., Ahanonu, B., Wu, Y.-W., Kim, T. H., Grewe, B. F., others (2018). Diametric neural ensemble dynamics in parkinsonian and dyskinetic states. Nature, 557(7704), 177–182.

Raz, A., Vaadia, E., & Bergman, H. (2000). Firing patterns and correlations of spontaneous discharge of pallidal neurons in the normal and the tremulous 1-methyl-4-phenyl-1, 2, 3, 6-tetrahydropyridine vervet model of parkinsonism. Journal of Neuroscience, 20(22), 8559–8571.

Redgrave, P., Prescott, T. J., & Gurney, K. (1999). The basal ganglia: a vertebrate solution to the selection problem? Neuroscience, 89(4), 1009–1023.

Reimann, M. W., King, J. G., Muller, E. B., Ramaswamy, S., & Markram, H. (2015). An algorithm to predict the connectome of neural microcircuits. Frontiers in Computational Neuroscience, 9, 120.

Reimann, M. W., Nolte, M., Scolamiero, M., Turner, K., Perin, R., Chindemi, G., Markram, H. (2017). Cliques of neurons bound into cavities provide a missing link between structure and function. Frontiers in computational neuroscience, 11, 48.

Salin, P., López, I. P., Kachidian, P., Barroso-Chinea, P., Rico, A. J., Gómez-Bautista, V., Lanciego, >J. L. (2009). Changes to interneuron-driven striatal microcircuits in a rat model of parkinson’s disease. Neurobiology of disease, 34(3), 545–552.

Sharott, A., Vinciati, F., Nakamura, K. C., & Magill, P. J. (2017). A population of indirect pathway striatal projection neurons is selectively entrained to parkinsonian beta oscillations. Journal of Neuroscience, 37(41), 9977–9998.

Shen, W., Zhai, S., & Surmeier, D. J. (2022). Striatal synaptic adaptations in parkinson’s disease. Neurobiology of Disease, 167, 105686.

Singh, A., Mewes, K., Gross, R. E., DeLong, M. R., Obeso, J. A., & Papa, S. M. (2016). Human striatal recordings reveal abnormal discharge of projection neurons in parkinson’s disease. Proceedings of the National Academy of Sciences, 113(34), 9629–9634.

Suarez, L. M., Solis, O., Sanz-Magro, A., Alberquilla, S., & Moratalla, R. (2020). Dopamine d1 receptors regulate spines in striatal direct-pathway and indirect-pathway neurons. Movement Disorders, 35(10), 1810–1821.

Surmeier, D. J., Graves, S. M., & Shen, W. (2014). Dopaminergic modulation of striatal networks in health and parkinson’s disease. Current opinion in neurobiology, 29, 109–117.

Tachibana, Y., Iwamuro, H., Kita, H., Takada, M., & Nambu, A. (2011). Subthalamo-pallidal interactions underlying parkinsonian neuronal oscillations in the primate basal ganglia. European Journal of Neuroscience, 34(9), 1470–1484.

Tanimura, A., Du, Y., Kondapalli, J., Wokosin, D. L., & Surmeier, D. J. (2019). Cholinergic interneurons amplify thalamostriatal excitation of striatal indirect pathway neurons in parkinson’s disease models. Neuron, 101(3), 444–458.

Taverna, S., Ilijic, E., & Surmeier, D. J. (2008). Recurrent collateral connections of striatal medium spiny neurons are disrupted in models of parkinson’s disease. Journal of Neuroscience, 28(21), 5504–5512.

Tsodyks, M. V., & Markram, H. (1997). The neural code between neocortical pyramidal neurons depends on neurotransmitter release probability. Proceedings of the National Academy of Sciences, 94(2), 719–723.

Underwood, C. F., & Parr-Brownlie, L. C. (2021). Primary motor cortex in parkinson’s disease: Functional changes and opportunities for neurostimulation. Neurobiology of Disease, 147, 105159.

Valsky, D., Heiman Grosberg, S., Israel, Z., Boraud, T., Bergman, H., & Deffains, M. (2020). What is the true discharge rate and pattern of the striatal projection neurons in parkinson’s disease and dystonia? Elife, 9, e57445.

Van Geit, W., Gevaert, M., Chindemi, G., Rössert, C., Courcol, J.-D., Muller, E. B., … Markram, H. (2016). Bluepyopt: leveraging open source software and cloud infrastructure to optimise model parameters in neuroscience. Frontiers in neuroinformatics, 10, 17.

Viaro, R., Morari, M., & Franchi, G. (2011). Progressive motor cortex functional reorganization following 6-hydroxydopamine lesioning in rats. Journal of Neuroscience, 31(12), 4544–4554.

Villalba, R. M., & Smith, Y. (2018). Loss and remodeling of striatal dendritic spines in parkinson’s disease: from homeostasis to maladaptive plasticity? Journal of neural transmission, 125, 431–447.

Wall, N. R., De La Parra, M., Callaway, E. M., & Kreitzer, A. C. (2013). Differential innervation of direct-and indirect-pathway striatal projection neurons. Neuron, 79(2), 347–360.

Zaja-Milatovic, S., Milatovic, D., Schantz, A., Zhang, J., Montine, K., Samii, A., … Montine, T. (2005). Dendritic degeneration in neostriatal medium spiny neurons in parkinson disease. Neurology, 64(3), 545–547.

Zhai, S., Shen, W., Graves, S. M., & Surmeier, D. J. (2019). Dopaminergic modulation of striatal function and parkinson’s disease. Journal of Neural Transmission, 126, 411–422.

Zhu, Y., Liu, B., Zheng, X., Wu, J., Chen, S., Chen, Z., … Lei, W. (2019). Partial decortication ameliorates dopamine depletion-induced striatal neuron lesions in rats. International Journal of Molecular Medicine, 44(4), 1414–1424.

Ztaou, S., & Amalric, M. (2019). Contribution of cholinergic interneurons to striatal pathophysiology in parkinson’s disease. Neurochemistry international, 126, 1–10.

