## Supplementary Material for "The impact of Parkinson’s disease on striatal network connectivity and cortico-striatal drive: an in-silico study"

*Journal:* NETWORK NEUROSCIENCE

RESEARCH

### SUPPLEMENTARY MATERIALS

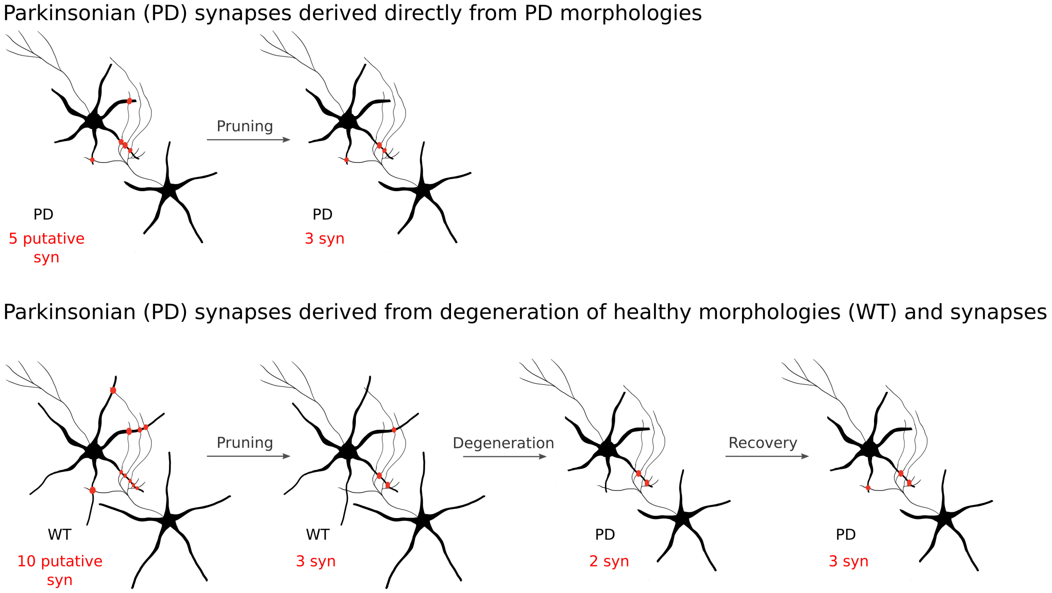


**Supplementary Figure 1: Determining the location of Parkinsonian (PD) synapses in the model network.** Because the pruning step will reduce the number of putative synapses to a narrow range, it will matter whether the pruning is done before or after the degeneration. In the first row of this example the (de novo) touch detection was done on the PD morphologies (after degeneration). Here 5 putative synapses were detected and three of those remain after the pruning step. In the second row the touch detection was done on the original WT (PD0) morphologies. Here due to the larger morphologies ten putative synapses were detected and the pruning reduces them down to three synapses. The subsequent degeneration then removes additional synapses on the lost dendrite branches, and only two synapses were left. However, we want a comparable final number of synapses as. Therefore a subset of the PD synapses from the de novo detection was added. This results in three PD synapses (in this example), one of which was recovered. These two different strategies give the same results with regard to whether a pair of neurons is coupled, and hence for the topological analysis. But in full microcircuit simulations the effects of the GABA interactions may be slightly different.

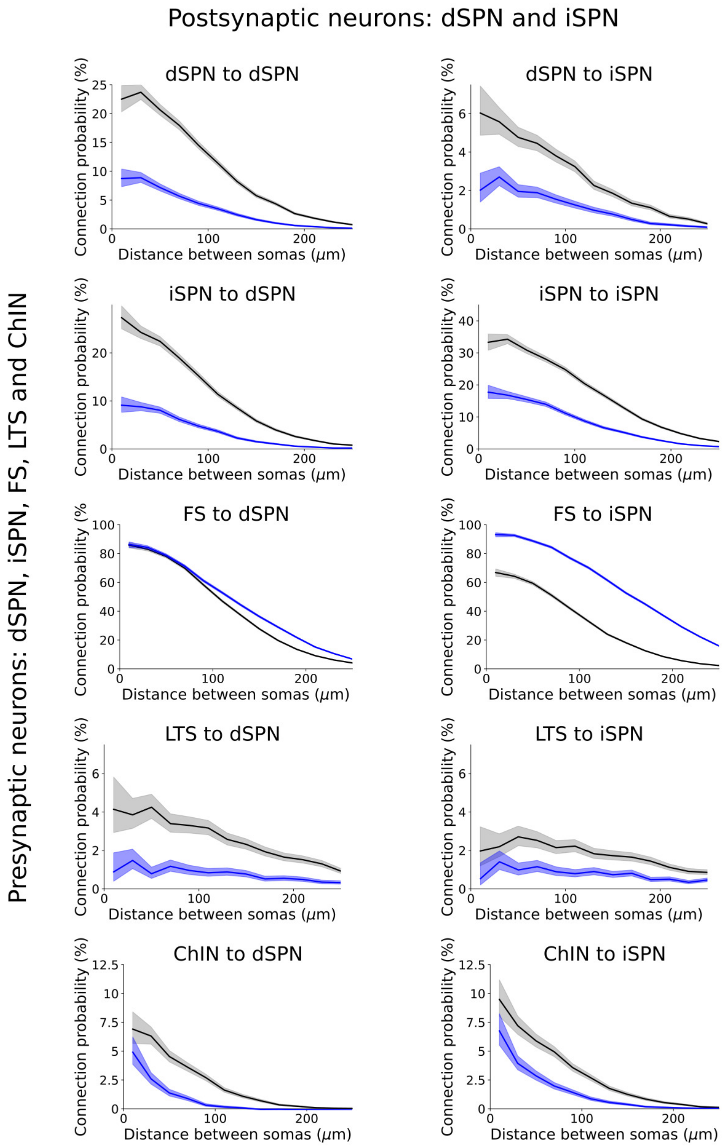


**Supplementary Figure 2**: **Pairwise connection probability at all soma-to-soma distances.**

In the first column there are pairwise connection probabilities at all soma to soma distances between the presynaptic neurons dSPN, iSPN, FS, LTS, ChiN and the postsynaptic dSPN. In the second column there are pairwise connection probabilities at all soma to soma distances between the presynaptic neurons dSPN, iSPN, FS, LTS, ChiN and the postsynaptic iSPN. Lines in black indicate the PD0 connection probabilities while in blue the PD2. The shaded regions represent the Wilson score interval.

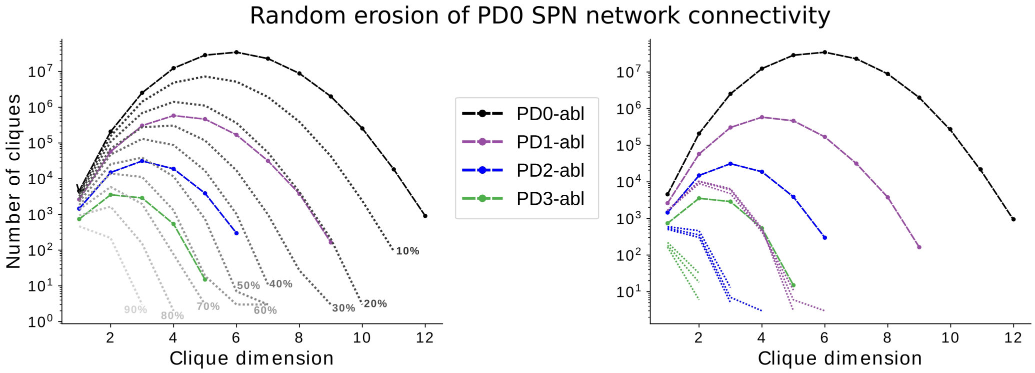


**Supplementary Figure 3: From healthy SPN striatal network to PD SPN networks.**

Number, in log scale, of cliques in PD0 (in black), PD1 (in purple), PD2 (in blue) and PD3 (in green) networks containing exclusively SPN neurons (interneurons are not included). Left: Random erosion of PD0 SPN-SPN synapses (between 90% and 10%) was performed and for each newly obtained network the directed clique distribution was plotted in a shade of grey. Right: Dotted distributions in purple, blue and green were obtained analysing eroded networks containing on average the same number of synapses as in PD1, PD2 and PD3 networks (where the SPN morphologies were degenerated). The removal of both proximal and distal synapses during the random erosion caused a more dramatic decrease in the both clique count and clique dimension compared to the distribution of directed cliques when only distal synapses were lost (because located on the the degenerated dendrites). This indicates that proximal synapses are more important for clique formation than distal ones.

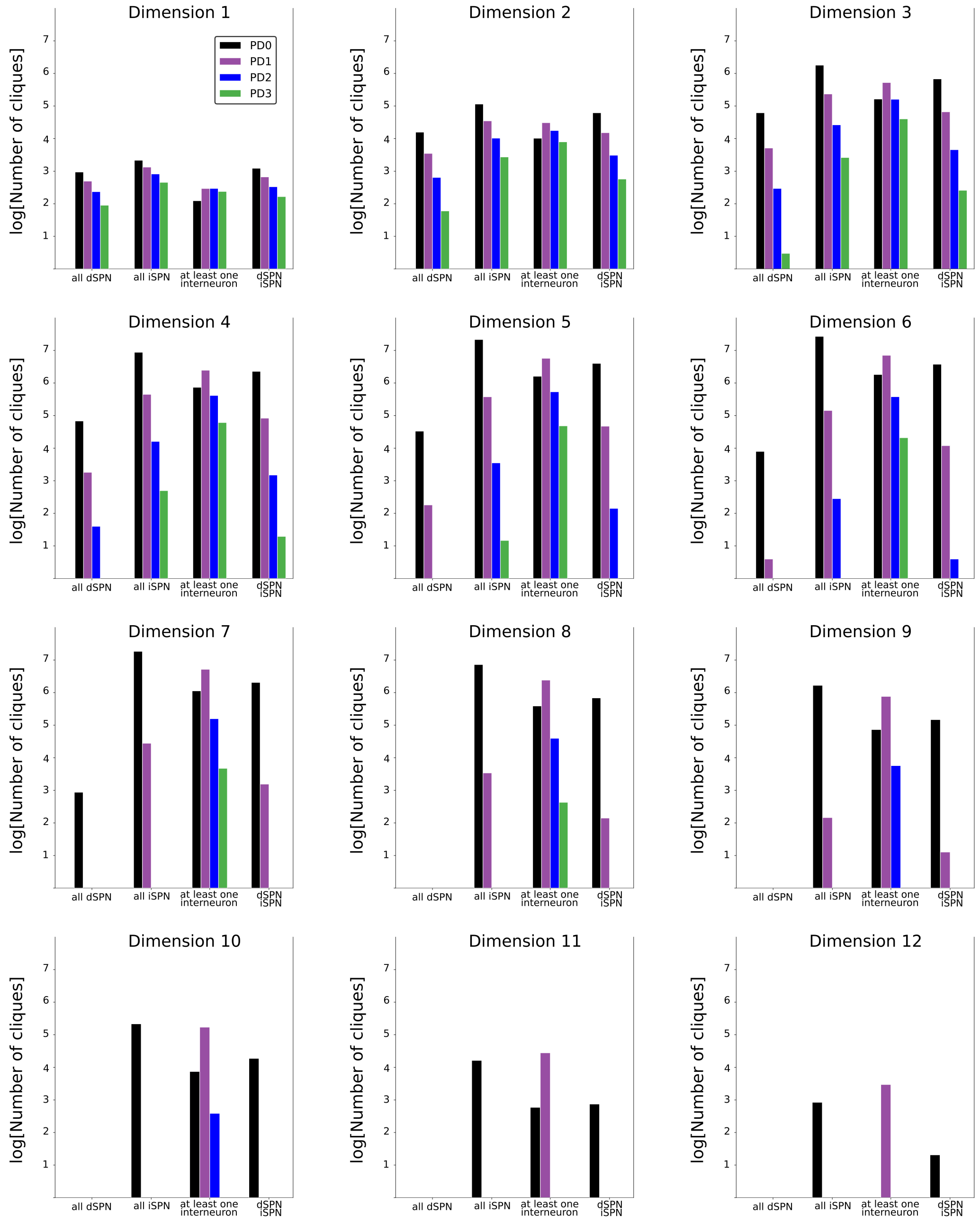


**Supplementary Figure 4**: **Clique composition.**

Number, in log scale, of cliques in PD0 (in black), PD1 (in purple), PD2 (in blue) and PD3 (in green) subdivided within the specific neuron compositions: all dSPN, all iSPN, at least one interneuron, both dSPN and iSPN.

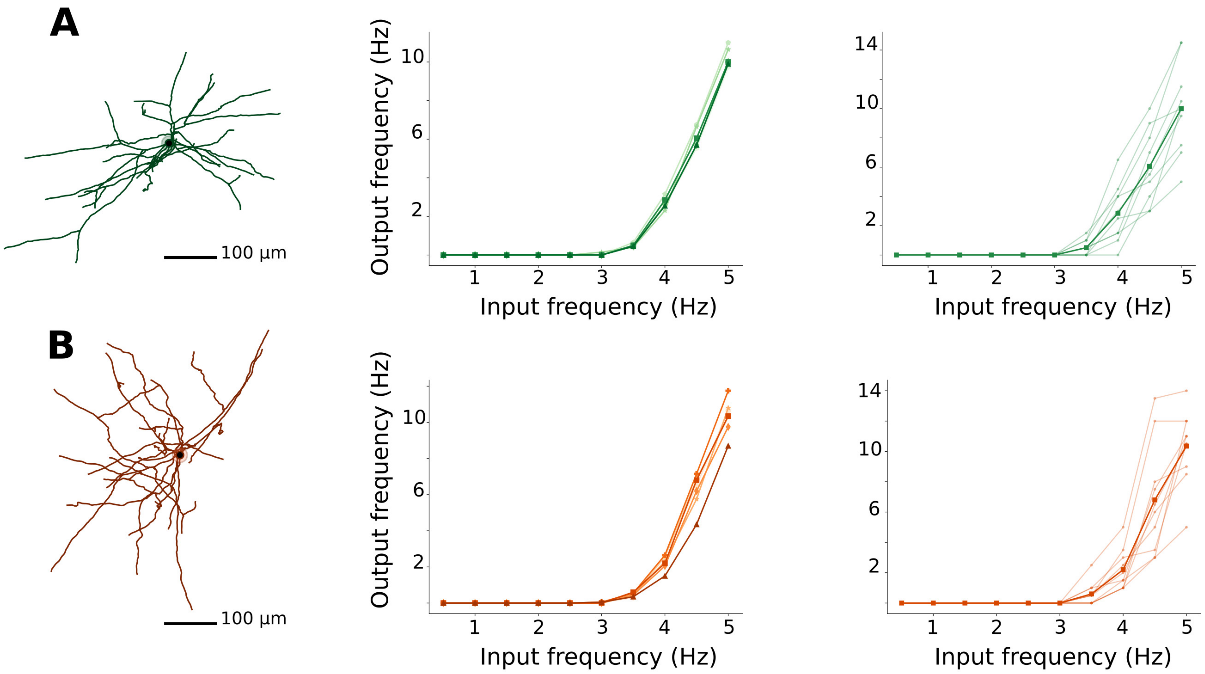


**Supplementary Figure 5**: **Tuning of the number of synapses in the healthy cells.**

Two variations of the same iSPN morphology are represented (see Materials and Method for details). The numbers of branches, stems and terminals are the same, but they differ for the total dendritic length, which corresponds to 3734 µm for the morphology in (A) and 4932 µm for the morphology in (B). As a consequence, the number of synapses needed to reach on average an output frequency of 10 Hz, when the activation frequency of each synapse is 5 Hz, differs. Around 170 and 210 synapses are required respectively. Each curve in the middle panels represents the average over 10 simulations of a specific (electric) parameters set. The same parameters sets are used both in A and B, but the morphology and the distribution of synapses vary. In the right panels an example of 10 such simulations are plotted. Despite having the same morphology, parameters set and number of synapses, the latter are distributed using different random seeds and this causes the variability in the response.

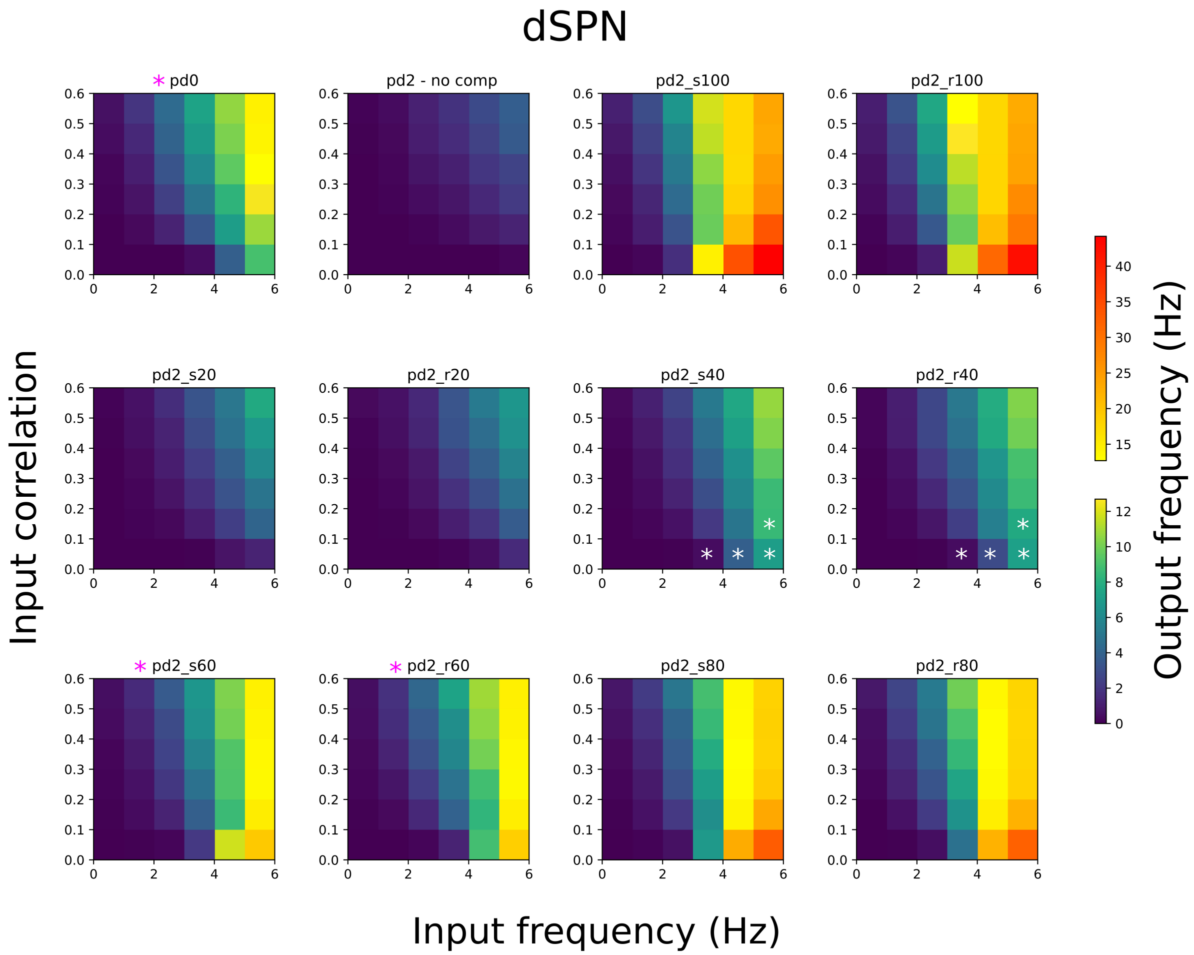


**Supplementary Figure 6**: **All combinations of input frequency, correlation and compensation in dSPN.**

Heatmaps summarising the relation between input frequency (x-axis), input correlation (y-axis) and output frequency (colormaps) in each stage. The healthy control case corresponds to "pd0", the pd without any compensation is denoted "pd2 - no comp", while the strengthening and rewiring is indicated as "pd2_s" and "pd2_r", respectively. For each compensation the numbers in the title represent the percentages of the compensations. Two colorbars are used in order to properly present the low output frequencies (in the blue-yellow colorbar). Moreover the low output frequency range is the same here and in the Supplementary Figure 6 where iSPN is presented. Purple asterisks indicate pd0 and the overall best approximations obtained with 60% compensation. White asterisks indicate the best approximation for a specific heatcell (input frequency-input correlation pair) when it does not belong to the overall best. It is interesting to notice that in some cases (pd2_s100, pd2_r100, pd2_s80, pd2_r80) some output frequencies at lower correlations are larger than those corresponding to higher correlations (see also Supplementary Figure 7). Averaging along the columns and rows of the heatmaps (pd2_s20, pd2_r20, pd2_s60, pd2_r60) gives the curves plotted in Figure 8B,C left and right panels respectively.

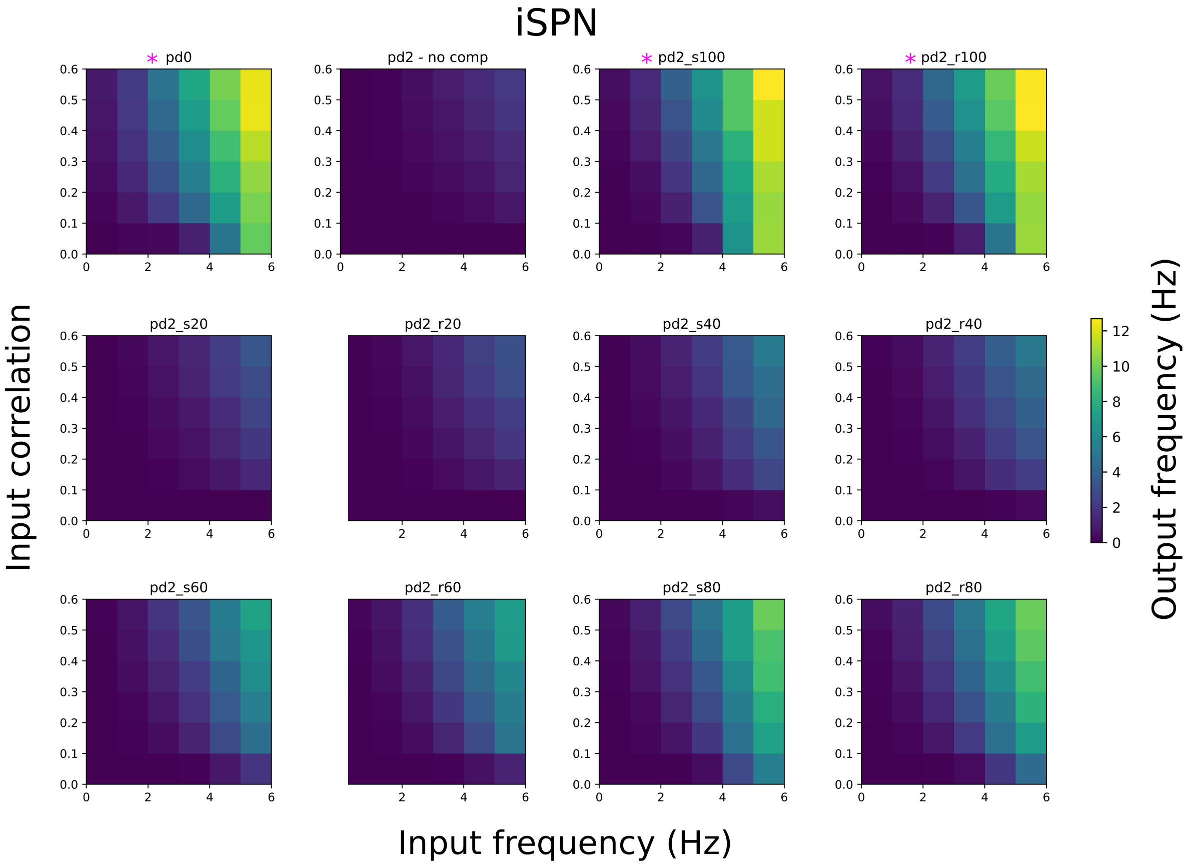


**Supplementary Figure 7**: **All combinations of input frequency, correlation and compensation in iSPN.**

Heatmaps summarising the relation between input frequency (x-axis), input correlation (y-axis) and output frequency (colormaps) in each stage. The healthy control case corresponds to "pd0", the pd without any compensation is denoted "pd2 - no comp", while the strengthening and rewiring is indicated as "pd2_s" and "pd2_r", respectively. For each compensation the numbers in the title represent the percentages of the compensations. The colorbar is used to present the output frequencies. Purple asterisks indicate pd0 and the overall best approximations obtained with 100% compensation. Averaging along the columns and rows of the heatmaps (pd2_s20, pd2_r20, pd2_s100, pd2_r100) gives the curves plotted in Figure 8D,E left and right panels respectively.

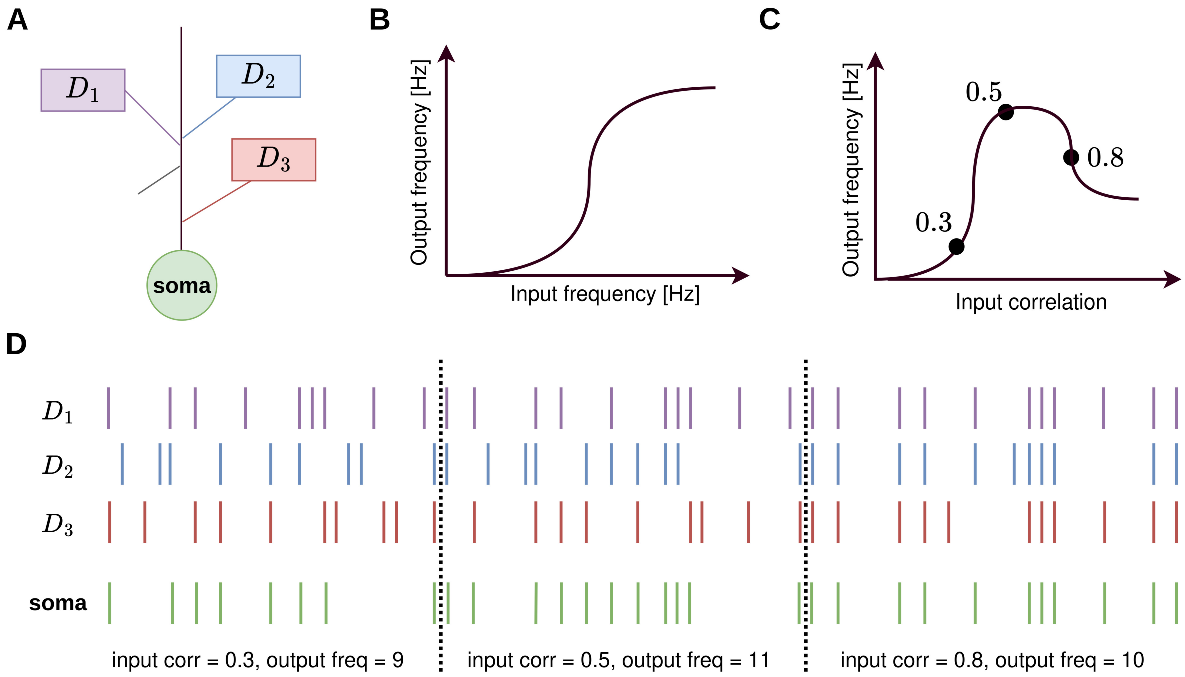


**Supplementary Figure 8: Input correlation versus output frequency - a conceptual explanation.**

A) The diagram depicts a schematic neuron with soma and dendritic branches, three of which are labelled D1 (purple), D2 (blue), and D3 (red). B) An example of an input-output function of a neuron, where output frequency increases with input frequency. C) A neuron's output frequency, given the same activation frequency of each synapse, is also influenced by the correlation between the input spikes, resulting in a non-monotonic input-output function. The three black dots correspond to different scenarios with regard to correlations. When all input spikes arrive at different times, the signal might be insufficient to activate the neuron. As input correlation increases, coincident spikes begin to activate the neuron, e.g., at an input correlation of 0.3. As input correlation continues to increase, the output firing rate also increases, e.g. at an input correlation of 0.5. However, when the input correlation becomes too high, input spikes are "wasted" as more "three-spike" coincidences appear in this simplified example, while only a "two-spike" coincidence is necessary to elicit an output spike, e.g. an input correlation of 0.8. D) Detailed illustration of how the neuron's input and output spikes under the varying input correlations in C. Each bar represents a spike, and each row represents a spike train where the colour corresponds to the label in A. For the three scenarios separated by dotted lines, the total input spikes remain constant, but the correlation changes. The neuron, here simplified as a coincidence detector, are assumed to spike whenever two or more coincident spikes arrive, as illustrated in the last row (green).

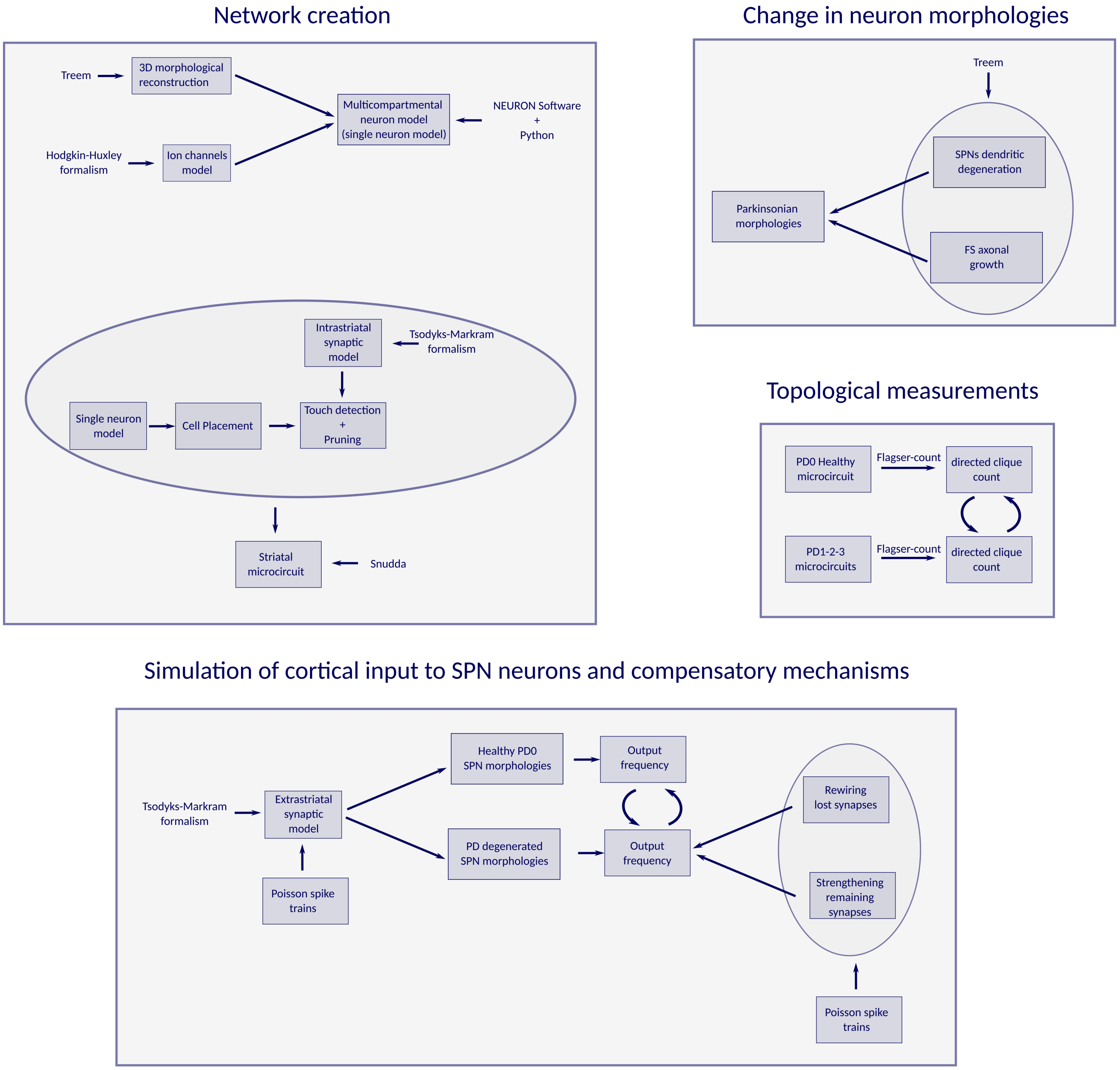


**Supplementary Figure 9**: **Box and arrow diagrams summarising the methods, software and tools used.** The diagrams summarise in a schematic way the steps of: network creation, change in neuron morphologies during PD, topological measurements, and simulation of cortical input to SPN neurons. This image is intended as support to better navigate the Materials and Methods section. Round arrows represent comparison analysis between the boxes they refer to.
